# Epigenomic circuits shaping cellular heterogeneity in pleural mesothelioma

**DOI:** 10.64898/2026.11.11.765995

**Authors:** Bruno Giotti, Lillian Chang, Komal Dolasia, Wooseung Lee, Subhasree Sridhar, Benjamin D. Hopkins, Amir Horowitz, Andrea Wolf, Yifei Sun, Anna Tocheva, Martin J. Walsh, Alexander M. Tsankov

## Abstract

Pleural mesothelioma (PM) is an aggressive and lethal malignancy with current treatments offering limited clinical benefit. In this study, we utilized single-cell chromatin accessibility and transcriptomic profiling of patient-matched samples to infer the epigenomic regulation underlying cellular heterogeneity in PM. High-resolution epigenomic profiling suggested transcriptional regulators with putative pro-tumoral roles in mesothelioma, whose function was experimentally tested using genetic perturbations across four PM cell lines. Furthermore, we observed several putative associations between epigenomic circuits and macrophage ontogeny, a sarcomatoid-like neoplastic state, and a fetal-like endothelial cell state. Our analysis also showed similarities between the chromatin accessibility landscapes of exhausted CD8 T and KLRC1 natural killer cells and identified NR4A2 as a potential shared regulator of dysfunction in both lymphocyte subsets. We further identified a state-specific distal enhancer near NR4A2 and experimentally validated its regulation of NR4A2 expression in Jurkat cells. Together, we provide an epigenomic map of PM and nominate candidate transcription factors and regulatory elements for future functional investigation, including their potential relevance to epigenetic editing strategies to limit tumor growth and enhance anti-tumor immunity.

## INTRODUCTION

Over the past decade, single-cell technologies have been instrumental in characterizing the cellular and molecular complexity of tumor microenvironments (TMEs) across human malignancies, often informing new therapeutic strategies and biomarkers (1). Similarly, Single-cell Assay for Transposase-Accessible Chromatin using sequencing (scATAC-seq) has shed light on the epigenomic circuits underpinning tumor transformation and plasticity of malignant cells (2–7). For example, scATAC-seq paired with single-cell RNA-sequencing (scRNA-seq) led to the discovery of a continuum of transcriptional programs governed by sequentially activated transcription factors (TFs) from healthy to cancer state via pre-cancerous polyps in colorectal cancer (CRC) (3) and revealed the regulatory logic driving tumor heterogeneity in gynecological cancers (6).

Characterizing the chromatin accessibility of TME cell types at single-cell resolution has also identified maladaptive regulatory circuits in cancer-associated stromal and immune cells (3,8–10). For instance, inferred TF activity from scATAC-seq implicated RUNX1 in regulation of cancer-associated fibroblasts (CAFs) in CRC (3) and identified multiple TFs driving the transition of naïve to dysfunctional CD8 T cells in basal cell carcinoma (8). Decoding the epigenomic rearrangements associated with lymphocyte dysfunction may inform strategies to improve the efficacy of current immune checkpoint inhibitors (ICI) therapies (11), and editing *cis*-regulatory elements (cREs) near immune checkpoint genes have shown improved persistence of the edited CD8 T cells compared to full PD-1-KO (12). Hence, deciphering the epigenome of tumor-infiltrated lymphocytes (TILs) has critical translational relevance.

Pleural mesothelioma (PM) is a rare and aggressive cancer arising from the pleura. Its pathogenesis is heavily linked to asbestos exposure and despite increased regulation in Western countries, PM world-wide incidence has not noticeably declined due to difficulties in environmental removal and lack of global regulation (13). Histologically, PM presents three main subtypes: epithelioid (50-60% of cases), sarcomatoid (10%), and biphasic (30-40%), which displays cell morphology of both epithelioid and sarcomatoid tumors (14). For many years, platinum-based chemotherapy has been the standard of care for the treatment of unresectable PM. Only recently have combination ICI treatments (nivolumab plus ipilimumab) demonstrated a significant enhancement in outcomes over chemotherapy, particularly for patients with non-epithelioid histology (15). Despite this, all subtypes of PM have an overall dismal prognosis and new therapeutic strategies are in dire need.

In the past ten years, genomic, transcriptomic and multiomic studies in PM have identified key genetic alterations, tumor subtypes, and TME molecular and cellular features of translational significance (16–21). However, no studies have investigated the chromatin accessibility landscape of PM at single-cell resolution preventing a deeper understanding of tumor cellular heterogeneity, identification of key regulators orchestrating immune-suppressive TME ecosystems, and discovery of novel biomarkers and therapeutic targets. Here, we provide a first effort in that direction, profiling the chromatin accessibility landscape of the PM TME using scATAC-seq, complemented by sample-matched scRNA-seq. Comparison of normal mesothelial and malignant cells identified multiple TFs with putative pro-tumoral roles in PM and further associated SOX9, SNAI2 and RUNX1/2 with a sarcomatoid-like neoplastic state. We also observed a putative regulatory dichotomy between monocyte-derived macrophages and tissue-resident macrophages, as well as previously underexplored similarities in the open chromatin landscapes of exhausted CD8 T cells and KLRC1 NK cells, both exhibiting high NR4A2 activity. Together, these analyses infer the regulatory circuits associated with cellular heterogeneity across various cell types in the PM TME, including immune-suppressive programs with potential implications for translational medicine.

## RESULTS

### Single-cell map of chromatin accessibility in PM

To explore the epigenome of PM at single-cell resolution, we performed high-throughput scATAC-seq on 11 treatment naïve tumors spanning all three PM histological types (**Supplementary Table 1**). We also profiled all but one sample from this cohort via scRNA-seq, which we used throughout the study to complement our epigenomic analyses (**Figure 1A**) (21). After thorough quality control (QC, **Methods**), we recovered 49,849 high-quality single-cell epigenomes (**Figure S1A-D**) and performed *de novo* cell annotation (**Methods**), capturing all expected PM cell types, including malignant, smooth muscle cells (SMCs), CAFs, vascular endothelial cells (VECs), mesothelial, alveolar, myeloid, T, B, plasma, natural killer (NK), and plasmacytoid dendritic cells (pDCs; **Figure 1B**). We detected a total of 401,152 unique chromatin accessibility peaks across cell types, where most peaks were identified in malignant cells (**Figure S1E**). Annotated cell clusters contained cells from multiple patient tumors except for malignant cells (**Figure 1C**, **S1F**), which formed patient-specific clusters likely due to differences in neoplastic genomic aberrations, as observed previously (7). TME cell classes showed highly specific patterns of chromatin accessibility enrichment at the gene locus, or genescore, of key marker genes (**Figure 1D**). To characterize the TF motif enrichments in open chromatin, we computed TF activity for each cell type using chromVAR (22) and identified the most differentially active TF motifs per cell type (**Figure 1E; Supplementary Table 2**). Predicted differential activities were consistent with established roles of several TFs in different cell types, including SOX13 in VECs (23), SPI1/SPIB in myeloid cells (24), MEF2C in SMCs (25), and TEAD1 in PM malignant cells (26).

**Figure 1.**
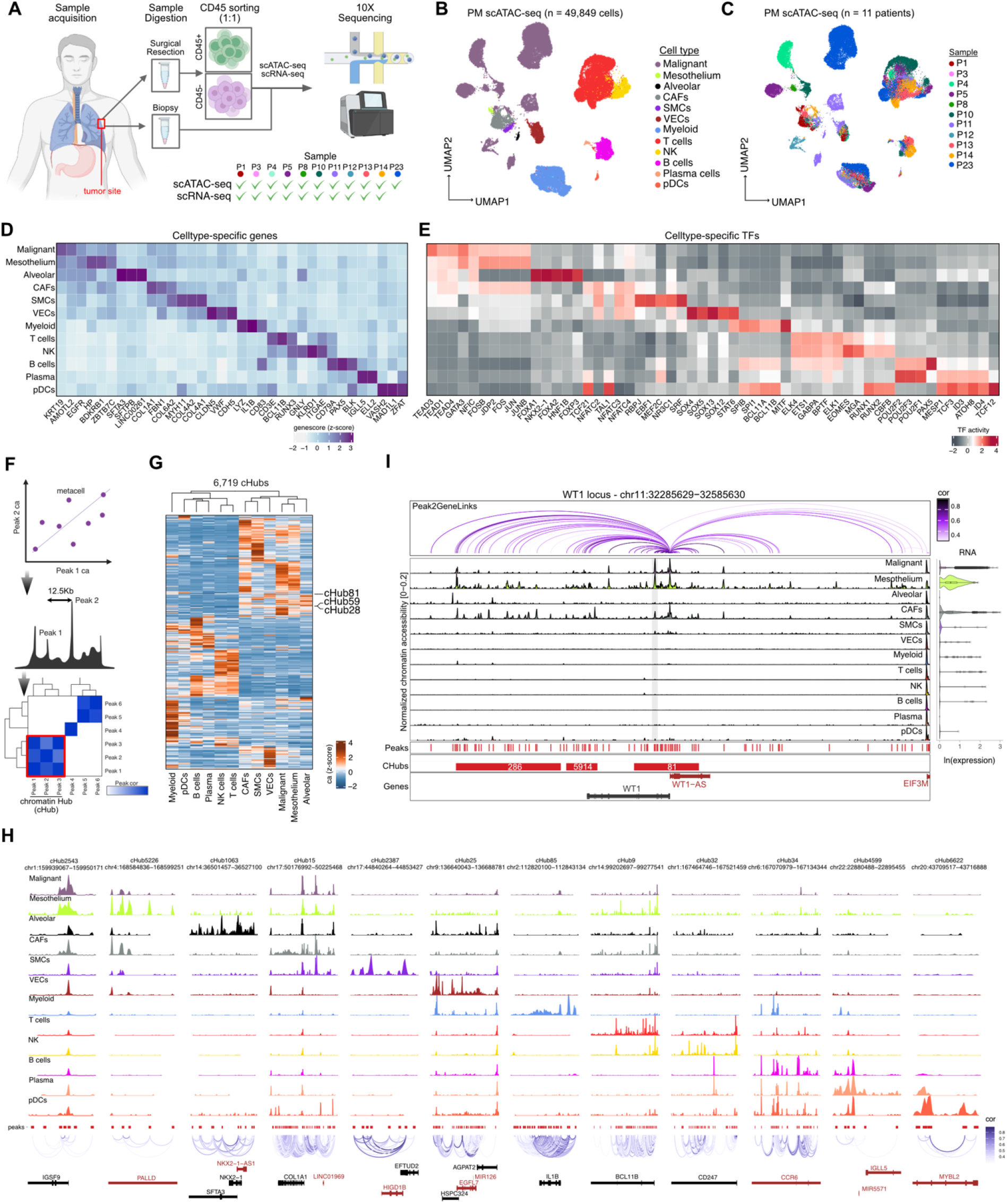
Single-cell atlas of chromatin accessibility in PM. **A,** Diagram of study’s experimental design. **B-C,** UMAP of integrated scATAC-seq data from 11 PM tumor samples colored by cell type annotation (**B**) and patient identity (**C**). **D,** Heatmap displaying the top three differentially enriched genes for each cell type based on genescore. **E,** Heatmap displaying the top five differentially active TFs for each cell type based on pre-computed chromVAR deviations. **F,** Schematic of the adopted algorithm used to detect cHubs. **G,** Heatmap showing the chromatin accessibility enrichment of all cHubs detected across cell types. Highlighted hubs were associated to WT1 gene. **H**, Examples of detected cHubs often associated with key lineage genes across annotated cell types. The number assigned to each cHub indicates its rank by region size in descending order (rank=1 denotes the largest). **I,** Pseudobulk chromatin accessibility tracks of all major cell types at the WT1 locus containing three cHubs (red bars). Also displayed are WT1’s RNA expression (violin plots, right), peak to gene correlation (purple arches, top), and highlighted region (grey vertical bar) shown in **Figure S2C**.

To confirm our malignant cell assignments, we complemented our marker gene annotations with copy number alteration (CNA) inference (**Methods**). Malignant cells from different tumors often displayed wide-spread chromosome losses compared to normal mesothelium, particularly on chromosomes 4, 13 and 22, in agreement with CNAs detected using patient-matched scRNA-seq data and previous CNA analysis from genomic data of larger PM cohorts (16,18,20) (**Figure S1G**). We further explored clonal heterogeneity within tumor samples using Epianeufinder (27) and identified subclones in five tumors (**Figure S1H**). Interestingly, we uncovered a subclone with amplification on the long arm of chromosome 8 (chr8q) in sample P4 that exhibited distinct spatial localization in four patient-matched tissue sections profiled by Visium spatial transcriptomics (**Figure S1I-J; Methods**). Finally, we asked which TFs showed the highest intra-tumoral variability in activity across the five tumors containing subclones and identified members of the AP1 complex and TEAD1, suggesting a convergent genetic selection for these TFs during subclonal evolution (**Figure S1K**).

Super-enhancers (SEs), or large transcriptionally active genomic regions, are frequently found upstream of lineage-determining genes in development and key oncogenes in cancer (28–30). To identify SE-like regions in PM, we adapted the Cicero algorithm (31) to identify clusters of open chromatin peaks with high co-accessibility across cells (**Figure 1F**, **Methods**), referred to as chromatin hubs (cHubs). Applying our approach to all scATAC-seq cells detected 6,719 cHubs (**Supplementary Table 3**), which showed highly unique patterns of chromatin accessibility across cell types (**Figure 1G**). The detected cHubs significantly overlapped SE regions in related primary cell lineages previously identified using bulk epigenomic data (28,32), validating the ability of our cHub analysis to detect cell type-specific SE-like regulatory regions from scATAC-seq data (**Figure S1L**, FDR-adjusted hypergeometric test). Furthermore, our cHub algorithm more accurately detected established SE regions compared to stitching nearby peaks as implemented in the ROSE algorithm (28), and was robust to different parameter settings (**Figure S1M-N**). Several cHubs were found nearby known key lineage genes, including COL1A1 in CAFs, SFTA3 and NKX2-1 in alveolar cells, and BCL11B in T cells (**Figure 1H**). Interestingly, we identified three cHubs at the locus of WT1, a key regulator of mesoderm and mesothelial development (33), displaying distinct chromatin accessibility landscapes in normal mesothelium, malignant cells, and CAFs (**Figure 1I**). To further characterize active DNA motifs at the WT1 locus, we trained a sequence-to-function deep learning model, chromBPNet (34) that predicted different TF motifs contributing to the locus chromatin accessibility in CAFs versus mesothelial cells, such as SNAI and ZN816/SMAD2, respectively (**Figures 1I, S2A-C**).

In summary, we present the first single-cell epigenomic map of PM, which enabled us to dissect the TME regulatory landscape at cellular resolution and catalogue cell type-specific cHubs.

### CRISPR perturbations of predicted PM pro-tumor TFs impact cancer cell proliferation

The ability of scATAC-seq data to accurately separate malignant and normal mesothelium cells allowed us to identify PM-associated TF regulatory activity at improved, cellular resolution. Dimensionality reduction of the 21,304 malignant cells displayed clustering of cells by patient identity, indicative of high inter-tumoral heterogeneity (**Figure 2A**). *De novo* clustering of malignant cells further highlighted intra-tumoral heterogeneity, identifying 21 clusters with differential enrichments of TF activity, including previously unreported members of the HOX gene family specifically enriched in tumor cluster C1 from sample P11 (**Figure 2B, Supplementary Table 2**). Pairwise correlation of TF activities across malignant cells identified TF activity clusters associated with hallmark cancer pathways, including epithelial to mesenchymal transition (EMT) (e.g., ZEB1, SNAI2, and TWIST1), interferon response (e.g., IRF3/7/9 and STAT2) and cell cycle (e.g., TFDP2 and E2F4) (**Figure S3A-B**). Finally, we identified TFs with consistently higher activity and RNA expression in neoplastic cells compared to normal mesothelial cells (35), revealing 68 TFs with putative pro-tumor function in PM (**Figure 2C**; **Supplementary Table 4**).

**Figure 2.**
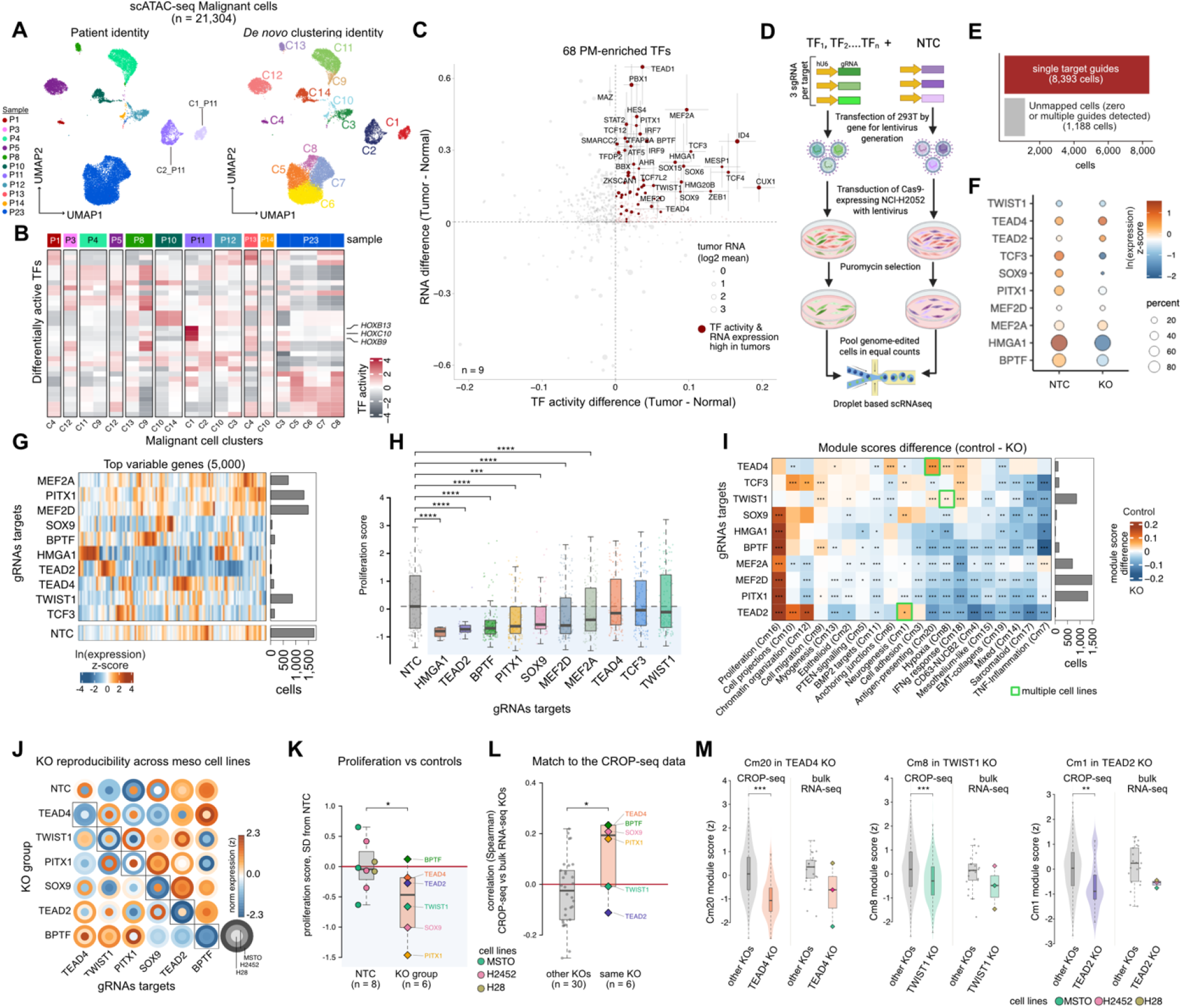
CRISPR droplet sequencing of computationally predicted pro-tumor TFs. **A,** UMAP of malignant cell scATAC-seq data color-coded by patient (left) and Louvain cluster identity (right). **B,** Heatmap displaying the most differentially active TFs across Louvain malignant cell clusters, where HOX family TF activities are specifically enriched in cluster C1. **C,** Scatter plot of the differences in median TF activity (x-axis) and median TF RNA expression (y-axis) between normal mesothelium and PM malignant cell samples identified 68 TFs (brown) with likely pro-tumor role. Error bars show standard error of the TF activity and RNA expression means in the tumor samples. **D,** Diagram of the CROP-seq experimental design. **E,** Majority of CROP-seq cells had single guides detected, while cells with zero or multiple guides detected were excluded from downstream analyses. **F,** Dot plot of the average expression and percentage of cells expressing TF targets in control and KO cells. **G,** Heatmap showing the pseudobulk expression profiles of aggregated control (bottom) and TF KO cells (top) across 5,000 most variable genes. Panel on right shows control and KO cell counts. **H,** Distribution of proliferation scores for control (grey) and KO (red) cells, Wilcoxon Rank Sum test, FDR adjusted. **I,** Differences in average module scores, calculated as control − KO; positive values indicate lower scores in KO cells. PM cancer modules were derived from (21). Wilcoxon Rank Sum test, FDR adjusted. **J,** Scaled expression of target genes across KO samples and NTCs for each of the three cell lines (concentric circles) with bulk RNA-seq data. **K,** Proliferation score of KOs relative to NTCs normalized as the standard deviation (SD) of KO sample scores from the mean NTC proliferation score per cell line (**Methods**). Circles, NTC samples colored by cell line; diamonds, KOs (median of SD from NTCs across cell lines); red line, mean NTC level. One-sided Wilcoxon rank-sum test, KO lower than NTCs. **L,** Spearman correlation between CROP-seq KO pseudobulk expression profiles and bulk RNA-seq KO expression profiles (averaged across cell lines) comparing correlation between corresponding KO pairs (colored) versus unmatched KO pairs (grey). **M,** Distribution of cancer module (Cm) scores in the targeted gene KO versus all other knockouts for Cm20 in TEAD4 vs other KOs, Cm8 in TWIST1 vs other KOs, and Cm1 in TEAD2 vs other KOs. Violin plots (left) compare CROP-seq data cells, while box plots (right) compare bulk RNA-seq samples averaged across cell line replicates. Two-sided Wilcoxon rank-sum tests was used to assess significance. Box plot horizontal lines show the 25th, 50th (median) and 75th percentiles, with vertical whiskers extending to a maximum distance of 1.5× the interquartile range from the hinge. Data beyond the whisker ends are plotted individually. *\*P* < 0.05, *\*\*P* < 0.01, *\*\*\*P* < 0.001, *\**\*\*\**P* < 0.0001, where *P* values > 0.05 are not shown.

To functionally test the role of the predicted pro-tumor TFs, we introduced Cas9-mediated genetic perturbations in the mesothelioma cell line NCI-H2052 via CRISPR droplet sequencing, or CROP-seq (36) (**Figure 2D**). Amongst the 68 TFs associated with high activity in PM, we selected 15 for targeting via CROP-seq, prioritizing TFs with detectable expression level in the NCI-H2052 cell line (RNA-seq data obtained from the Cancer Cell Line Encyclopedia (CCLE) (37)), whose RNA expression is associated with poor overall survival (16,17), and with likely compensatory function within TF family (e.g., TEAD, MEF) members (**Figure S3C**). After processing of the CROP-seq data (**Methods**), we recovered 8,393 high-quality scRNA-seq cells containing a unique guide RNA (gRNA) target for 10 of the 15 TFs screened, whereas 1,188 cells detected zero or multiple gRNA targets and were thus discarded from further analysis (**Figures 2E, S3D**). As additional QC, we validated the cell line identity of the CROP-seq data by comparing each cell’s CNA landscape, inferred using the scRNA-seq data, with CNA patterns derived from CCLE whole-exome and bulk RNA sequencing data for cell line NCI-H2052 (37), which were largely in agreement (**Figure S3E, Methods**). Expectedly, gRNAs targeting the same genes showed similar pseudo-bulk expression profiles and were grouped together for downstream analyses (**Figure S3F**). To evaluate the efficacy of the mediated individual perturbations, we first compared the TFs’ expression in knockout (KO) versus cells that received non-targeting control gRNAs (NTC), which showed a significant decrease in expression for TCF3, SOX9, HMGA1, PITX1, and BPTF (**Figure 2F, S3G,** Wilcoxon rank-sum test). As gene editing may leave target gene expression levels unchanged while generating non-functional transcripts, we further examined the top 5,000 variable genes across KO groups identifying clear differences in their pseudo-bulk transcriptional profiles compared to control cells, suggesting effective CRISPR editing in the remaining KOs (**Figures 2G**).

To assess the functional role of each TF perturbation, we first computed a cell cycle score for all cells in the CROP-seq data and found reduced cell proliferation for all 10 TF KOs compared to control cells, which was significant for 7 of 10 KOs (**Figure 2H**, Wilcoxon Rank-Sum Test, FDR adjusted). To dissect the functional effects of the TF KOs beyond cell proliferation, we quantified the per cell activity of 20 PM malignant programs we previously defined using scRNA-seq of 13 PM patient tumors (21) and computed the difference in program scores between control and KO cells (**Figure 2I**, Wilcoxon rank-sum test, FDR adjusted). This analysis identified unique changes in cancer programs expression for different TF KOs. For example, TWIST1 KO showed depletion in gene modules linked to cell migration and hypoxia (Cm9, Cm8, respectively), in line with prior studies showing that TWIST1 triggers EMT and is a direct target of hypoxia-inducible factor-1alpha (HIF-1alpha) in human cell lines (38). We also report on distinct transcriptional pathways affected by perturbation of TEAD members as TEAD2 showed downregulation of neurogenesis pathways (Cm1), consistent with its involvement in regulating neural progenitors with YAP (39), whereas TEAD4 KO induced down-regulation of antigen-presenting module (Cm20), which is consistent with its known role in Hippo pathway dysregulation (40) and previous literature linking hippo-signaling and MHC-II antigen presentation in tumor cells (41).

To test the generalizability of our CROP-seq results, we performed CRISPR KOs in the same manner for 6 of 10 TFs in three additional mesothelioma cell lines followed by bulk RNA-seq profiling. In total, 43 RNA-seq samples were of high quality as assessed by read coverage, matching cell line CNA profiles with corresponding CCLE lines, presence of gRNA reads in corresponding KO samples, high correlation of biological replicates (**Figure S4A-D,** one-sided Wilcoxon rank-sum test, FDR adjusted), and reduced expression of target TFs that was largely consistent across cell lines (**Figure 2J)**. We observed a similar lower proliferation trend in most of the KOs compared to NTCs (**Figure 2K, S4E,** one-sided Wilcoxon rank-sum test) as well as an overall agreement in the KO transcriptional profiles of the different cell lines with the CROP-seq profiled cell line (**Figure 2L, S4F,** one-sided Wilcoxon rank-sum test). Lastly, we confirmed KO trends in at least two of the three additional cell lines tested for Cm20 in TEAD4 KO, Cm8 in TWIST1 KO, and Cm1 in TEAD2 KO (**Figure 2M, 2I)**.

Overall, we systematically identified 68 putative pro-tumor TFs in PM and functionally tested the impact of a subset of these TFs on cell proliferation and other cancer-intrinsic expression programs with congruent results across multiple mesothelioma cell lines, demonstrating the predictive value of our PM scATAC-seq atlas.

### SOX9, SNAI2, and RUNX2 activity associates with sarcomatoid-like malignant cell state

Sarcomatoid PM tumors are more resistant to a wide range of chemotherapies relative to epithelioid PM (42); hence, high-resolution dissection of the regulatory drivers of sarcomatoid neoplastic state could inform on novel strategies aimed at eradicating this more aggressive and refractory PM subtype (42). To do so, we first scored malignant cells based on a sarcomatoid gene set (scS-score; **Supplementary Table 5**) we derived previously (21) and compared them with chromatin accessibility profiles of normal cell types in lung tissue (35) (**Figure 3A-B**; **Methods**). As expected, malignant cells with high scS-score were most correlated with the open chromatin landscape of normal lung mesenchymal cells, including fibroblasts and SMCs, and anti-correlated with epithelial cells (e.g., alveolar type 1 (AT1) and type 2 (AT2), secretory, ciliated) chromatin accessibility (**Figure 3B**). Correlating the scS-score with a large compendium of histone H3 lysine 27 acetyl (H3K27Ac) chromatin immunoprecipitation and sequencing (ChIP-seq) data from ENCODE as well as *in vitro* differentiated and primary cells showed concordant results (**Figure S5A**). For example, the epigenomes of fibroblast-derived cell lines (e.g., IMR-90, SJSA1) and fibroblast primary cells from foreskin, dermis and lung were mostly correlated with scS-score, whereas colon adenocarcinoma cell line Caco-2 and *in vitro* differentiated “mesothelial cell of epicardium” were most anti-correlated (**Figure S5A**).

**Figure 3.**
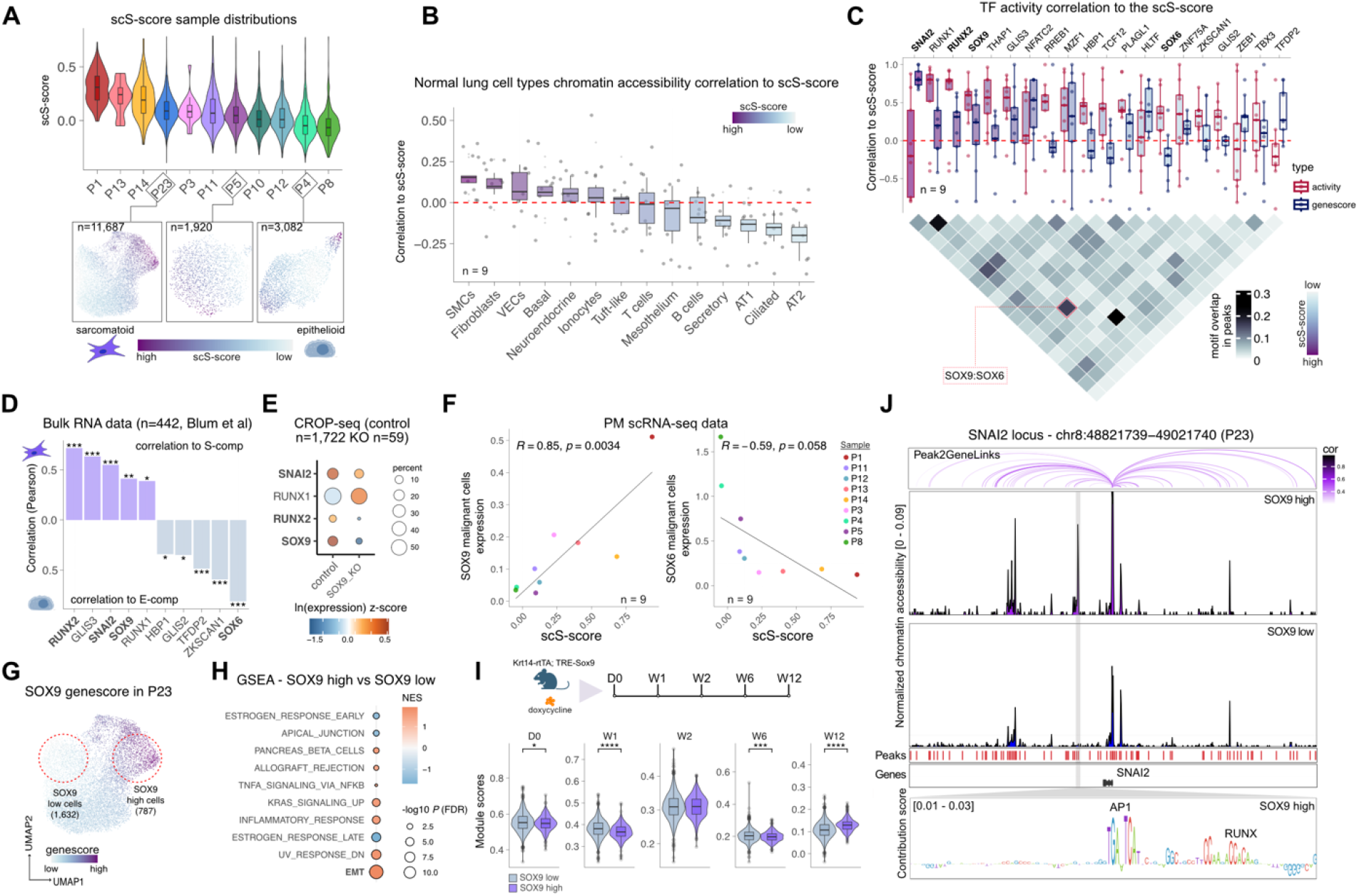
SOX9, RUNX2, and SNAI2 activity associates with sarcomatoid-like neoplastic state. **A,** Malignant cell distribution of scS-scores (top) for each PM tumor sample and UMAPs of three highlighted biphasic tumor samples malignant cells colored by their scS-score (bottom). The scS-score was computed per malignant cell based on a PM sarcomatoid gene set defined previously (21) (**Methods)**. **B,** Sample-level distributions of Spearman correlations between malignant cells’ scS-score and accessibility of open chromatin sets of peaks for different normal lung cell types (35). **C,** Top: Sample-level distribution of Spearman correlations between malignant cells’ scS-score and TF activity (red) or TF genescore (blue). Bottom: Pairwise TF motif overlap shows a high overlap of SOX9 and SOX6 motifs in malignant cell peaks. **D,** Correlation across bulk RNA-seq samples between TF (from **C**) expression and S-comp (sarcomatoid score) or E-comp (epithelioid score) derived from a meta-analysis study on PM (19). Pearson correlation *P* values. **E,** CROP-seq data expression and percentage cells expressing for TFs SNAI2, RUNX1, RUNX2, and SOX9 in SOX9 KO versus control cells. **F,** Scatter plots showing the average scS-score in malignant cells (x-axis) and the average expression of TFs (y-axes) SOX9 (left) and SOX6 (right) across PM scRNA-seq samples. Spearman rank correlation coefficient and *P* value are displayed on top. **G,** UMAP of PM sample P23 malignant cells colored by SOX9 genescore demonstrating assignment of SOX9 high and SOX9 low cells. **H,** GSEA enrichment analysis showing the most enriched HALLMARK gene sets for differentially accessible genes in SOX9 high (red) and SOX9 low (blue) cells. **I,** Distribution of module scores for gene sets found to be up-regulated at different time-points in SOX9 low and high cells—day 0 (D0), week 1 (W1), week 2 (W2), week 6 (W6), and week 12 (W12)—after induction of a Sox9 transgene in murine epidermal stem cells (43) in SOX9 high (purple) and SOX9 low (blue) cells. Wilcoxon Rank Sum test. **J,** Genome browser of the SNAI2 locus highlights an open chromatin region specific to SOX9 high cells (grey vertical bar) harboring a RUNX binding site predicted to be active using chromBPNet (bottom). Also displayed peak to gene correlation (purple arches, top). Box plot horizontal lines show the 25th, 50th (median) and 75th percentiles, with vertical whiskers extending to a maximum distance of 1.5× the interquartile range from the hinge. Data beyond the whisker ends are plotted individually. \**P* < 0.05, \*\**P* < 0.01, \*\*\**P* < 0.001, \*\*\*\**P* < 0.0001, where *P* values > 0.05 are not shown.

Next, we identified TFs whose activities/genescores were most correlated with scS-score to uncover putative regulators of PM sarcomatoid-like state and found SOX9, RUNX1/2, and SNAI2 among the top inferred TFs (**Figure 3C, top**). In agreement, expression of SOX9, RUNX1/2, and SNAI2 was also highly correlated with a previously derived sarcomatoid signature (S-comp) across a compendium of bulk RNA data from multiple independent PM cohorts (19) (**Figure 3D**). In animal models, Sox9 has been shown to directly induce Runx1/2 TFs (43) and to be bound at the *Snai2* promoter during neural crest formation (44). Consistently, RUNX2 and SNAI2 were downregulated in SOX9 KO versus control cells from our CROP-seq experiment, suggesting that SOX9 may also induce RUNX2 and SNAI2 expression in PM cell line NCI-H2052 (**Figure 3E**). In contrast, we noticed high correlation of sarcomatoid score with SOX6 TF activity, but anti-correlation with SOX6 genescore and RNA expression in PM tumors (**Figure 3C-D, 3F**). Since SOX6 shares a similar DNA binding motif with SOX9 (**Figure 3C**, **bottom**; **S5B**), it is possible that these two TFs compete for similar binding sites that may drive PM malignant cells to opposing ends of the scS-score gradient. In agreement, the association of SOX9, RUNX2, SNAI2, and SOX6 expression with the sarcomatoid-epithelioid histological and molecular subtypes was highly concordant across bulk RNA-seq data from three independent PM cohorts (**Figure S5C**). Since sarcomatoid PM is historically associated with worse survival, we next asked if the expression of the three sarcomatoid-associated TFs was linked to poor outcome in the same bulk RNA-seq cohorts. Indeed, we found SOX9 expression to be associated with worse overall survival, even after adjusting for sarcomatoid content in the Bueno et al. cohort (adjusted *P* = 0.009, Cox proportional-hazards model; **Figure S5D**), suggesting a prognostic value for SOX9 expression in PM disease outcome.

To further investigate the role of SOX9 in intratumoral cell heterogeneity, we focused on sample P23 that contained the highest number of SOX9 high (n=787) and SOX9 low (n=1,632) cells (**Figure 3G)**. As expected, SOX9 high/low cell status positively correlated with RUNX2 and SNAI2 genescores (**Figure S5E**). Gene set enrichment analysis (GSEA) on differentially accessible genes (DAG) in SOX9 high versus low cells demonstrated high enrichment for EMT and depletion for apical junction pathways in SOX9 high cells, consistent with increased mesenchymal properties and loss of epithelial function (**Figure 3H).** SOX9 high cells also showed significantly higher genescore for genes up-regulated at 12 weeks following inducible expression of a *Sox9* transgene in murine epidermal stem cells (43), further supporting a link with SOX9-induced gene programs (**Figure 3I**). Training chromBPNet models in SOX9 high and low cells highlighted SNAI among the most active TF motifs in SOX9 high cells (**Figure S5F-G).** Finally, we examined the chromatin accessibility at the SNAI2 locus and identified a distal peak specific to SOX9 high cells that included a RUNX motif, further pointing at an association between SOX9, RUNX2, and SNAI2 (**Figure 3J**).

Taken together, our scATAC-seq data identified TFs SOX9, RUNX2, and SNAI2 as likely regulators of sarcomatoid-like malignant cells and suggest a feed-forward regulatory circuitry amongst these factors that may reinforce their activity.

### Epigenetic-based regulatory dichotomy associated with putative macrophage ontogeny

Tumor-associated macrophages (TAMs) regulate cytotoxic lymphocytes, hamper efficacy of ICI treatments and correlate with worse overall survival in PM (45,46). We annotated 6,394 scATAC-seq myeloid cells, which broadly partitioned into monocytes, dendritic cells (DCs), and TAMs based on genecores of established markers (**Figures 4A, S6A-B**). To further dissect myeloid cell heterogeneity, we performed *de novo* gene module discovery using consensus non-negative matrix factorization (cNMF) on our sample-matched scRNA-seq data (**Methods**). Via cNMF, we identified 10 myeloid gene modules conserved across samples, where four closely resembled previously characterized TAM-associated cell subsets (47): TAM MARCO (module.1), *CXCL9/10/11* expressing TAM CXCLs (module.3), TAM interstitial (module.10), and TAM TREM2 (module.7). We also identified two monocyte modules, Mono CD16 (module.2) and Mono CD14 (module.6), as well as a DC (module.5), a cell cycle (module.4), a stress-related module (module.9), and a TAM module expressing CD52 (module.8) that lacked a coherent signature across samples and was not used in downstream analyses (**Figure S6C**; **Supplementary Table 6**). We used these modules to score myeloid cells in our scATAC-seq cohort and assigned each cell to a module via K-means clustering, excluding module.8 as well as the cell cycle and stress modules (module.4 and module.9), since they do not represent myeloid-specific expression programs (**Figure 4B**). These annotations enabled identification of TF motifs specifically enriched in the seven myeloid cell subsets. For example, we found higher activity of CCAAT/enhancer-binding protein (CEBP) TF motif in monocytes (48). Additionally, Erythroblast Transformation Specific (ETS) TFs family, known to play important roles in immune cell and macrophage development (49,50), including SPI1 (encoding PU.1), SPIB, ELF1, and ETV6, demonstrated elevated motif activity in TAM interstitial subset (**Figure 4C**; **Supplementary Table 2**).

**Figure 4.**
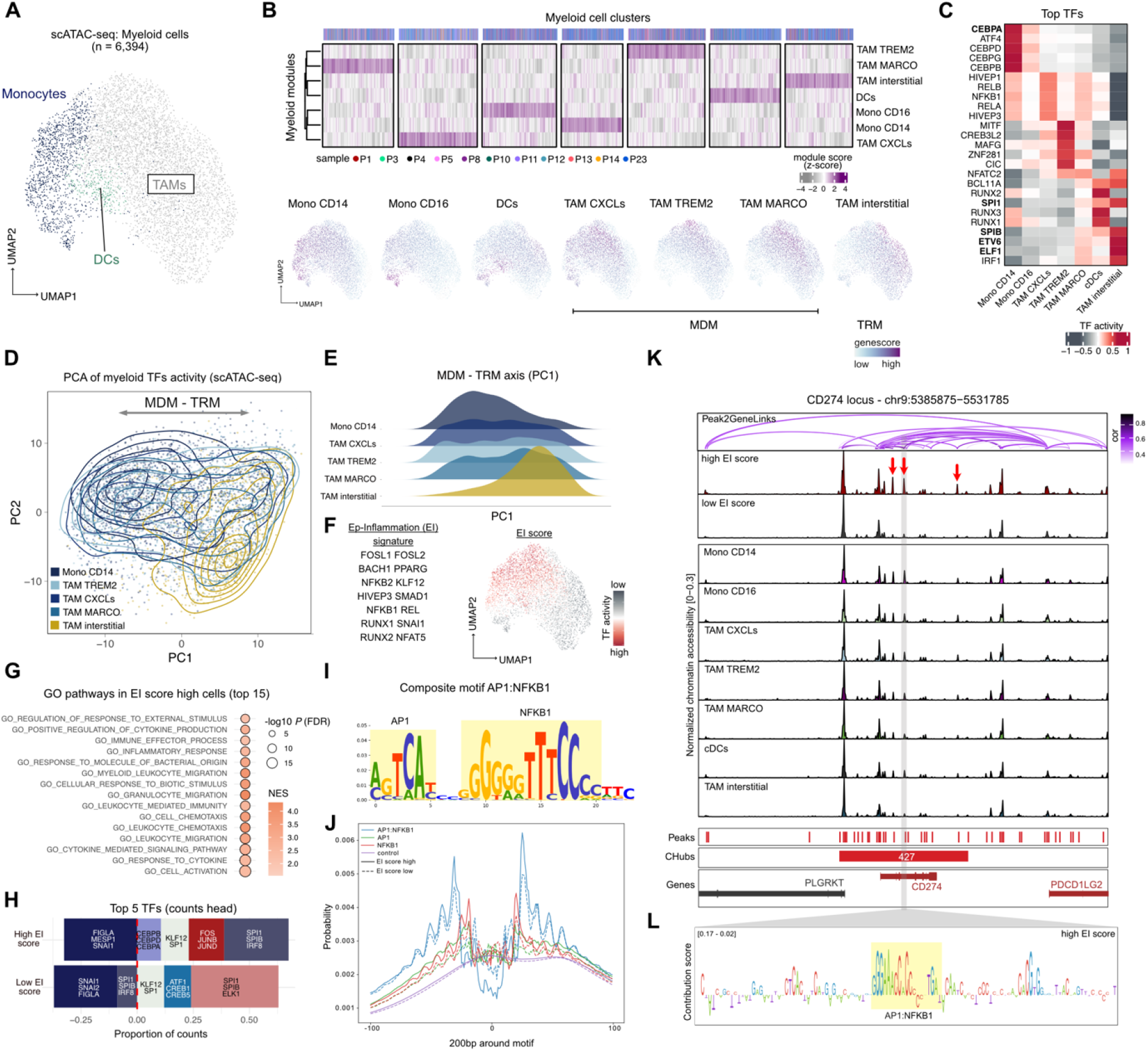
Epigenetic-based regulatory dichotomy captures putative macrophage ontogeny. **A,** UMAP of myeloid cell scATAC-seq data colored by broad cell types. **B,** Top: Heatmap showing 7 myeloid cell clusters (top) and their normalized score for 7 myeloid modules (right) derived from scRNA-seq paired samples, defining TAM subsets, DCs, and monocytes across all scATAC-seq myeloid cells. Bottom: UMAPs of myeloid cells colored by the normalized score for each of the 7 myeloid modules. **C,** Heatmap displaying the top 5 differentially active TFs in annotated myeloid subsets. **D,** PCA plot showing PC2 versus PC1 scores calculated using all TF activities in Mono CD14 and TAM subsets. PC1 captured the MDM-TRM epigenomic regulatory axis. Contours depict the density of different myeloid subsets. **E,** Distribution of PC1 scores for each myeloid subset. **F,** Left: List of top 14 TFs associated with PC1 (MDM-TRM axis) and used to compute the EI score for all myeloid cells (**Methods)**. Right: UMAP from panel **A** colored by EI score. **G,** GSEA analysis showing the 15 GO ontology pathways most enriched for DAGs between high versus low EI score cells. **H,** Top 5 most frequent motifs calls detected from chromBPNet counts outputs after training chromBPNet models using high and low EI score cells. Motifs with negative proportions are associated with negative contributions to chromatin accessibility. **I,** Composite AP1:NFKB1 motif identified by TF-MoDISco on chromBPNet-generated contribution scores. **J,** Footprints of AP1, NFKB1, and composite AP1:NFKB1 (from **I**) motifs in chromBPNet models trained on high and low EI score cells**. K**, Genome browser of the CD274 locus shows pseudobulk chromatin accessibility for high and low EI score cells as well as all myeloid subsets. Also shown are regions with differential chromatin accessibility across myeloid subsets (red arrows), peak to gene correlation (purple arches, top), and detected cHubs (red bars, bottom). Number assigned to each cHub indicates its rank by region size with 1 denoting the largest, where cHubs were recomputed within myeloid cells. **L,** Instance of AP-1:NFKB1 composite motif within the highlighted differentially accessible region (grey) at the CD274 locus.

Since monocyte-derived macrophages (MDM) are developmentally distinct from tissue-resident macrophages (TRM), we wanted to investigate if their chromatin accessibility profiles in PM may reflect this lineage ontogeny. To do this, we performed principal component analysis (PCA) on the per cell TF activities of Mono CD14 and TAM cell subsets (**Figure 4D**). We found that the first principal component (PC1) identified a regulatory gradient from Mono CD14 to MDMs to TRMs (TAM interstitial), demarcating a putative MDM-TRM axis and suggesting that TF activity primarily explained differences in macrophage ontogeny (**Figure 4D-E).** Interestingly, when we applied a similar analysis on sample-matched scRNA-seq data using TF regulon gene expression as a surrogate for TF activity (**Methods)**, PC1 now captured the variation between monocytes and all TAM subsets, irrespective of myeloid cell ontogeny (**Figure S6D-E**). The putative MDM-TRM regulatory axis was not captured by PC2 (driven by cell cycle, not shown) or any other PCs in RNA expression space (**Figure S6F)**. We observed similar trends using Multiome data (joint scRNA-seq and scATAC-seq on the same cells) from non-small cell lung cancer (NSCLC) derived myeloid cells (10) (**Figure S6G-H**). Moreover, co-activity of TFs across myeloid cells within PM and NSCLC datasets independently supported two distinct regulatory networks that tracked with macrophage ontogeny (**Figure S6I**).

We then sought to better characterize the putative MDM-TRM regulatory axis by inspecting the TFs most associated with it. Instead of myeloid lineage TFs, we found cell stress and inflammatory-related TF motif activities as most correlated with the MDM-TRM axis, including the AP-1 complex members FOSL1/2 and BACH1 and members of the nuclear factor-kappa B complex NFKB1/2 and REL, which we combined to compute a per cell average activity that we named the Ep-Inflammation (EI) score (**Figure 4F, S6J, Supplementary Table 5**). To explore biological differences between cells with high and low EI score, we performed GSEA on DAGs between these two groups of cells and expectedly identified Gene Ontology (GO) pathways (51) related to cytokine signaling, chemotaxis, and inflammatory response associated with high EI score cells (**Figure 4G**). We then trained chromBPNet models for high and low EI score cells, which similarly identified a higher fraction of the AP1 complex motifs (FOS/JUNB/JUND) in high EI score cells, as well as monocyte regulators CEBPB/D/A (**Figure 4H**). Among the *de novo* motifs identified by chromBPNet, we identified a composite motif comprising AP-1 and NFKB1 binding motifs that was associated with a more prominent footprint compared to AP1 or NFKB1 binding sites alone (**Figure 4J).** To further investigate the potential link between EI signature and putative macrophage ontogeny in an external dataset, we used published normal human Peripheral Blood Mononuclear Cells (PBMCs) and Bone Marrow Mononuclear Cells (BMMCs) scATAC-seq data (52) as a reference set to label our PM myeloid cells. Consistent with a putative MDM-TRM axis, PM monocyte and MDM cell subsets exhibited higher similarity to PBMC monocytes than PM TRMs (**Figure S6K-L**).

To identify large genomic open chromatin domains in myeloid cells, we used our cHub algorithm to detect 2,271 myeloid-specific cHubs (**Supplementary Table 3**, **Methods**), including several differentially accessed cHubs (DAcH) at the loci of key myeloid markers, such as IL1A-IL1B in monocytes and complement C1Q-associated subunits in TAM MARCO subset cells (**Figure S6M**). Similarly, we identified several DAcH between cells with high and low EI score, including cHub.427 at the CD274 (encoding PD-L1) gene locus (**Figure 4K, S6N**). This locus displayed elevated levels of chromatin accessibility in high EI score cells (**Figure 4K**, red arrows) and contained an instance of the composite AP1:NFKB1 at the highlighted region (**Figure 4L**), both suggestive of EI-mediated regulation of CD274 in a subset of myeloid cells.

In sum, we dissected myeloid epigenomic heterogeneity in PM and identified an EI regulatory cluster enriched for NF-κB and AP-1 complex TFs motifs and reflective of a putative ontogeny-associated regulatory axis in PM and NSCLC tumors. Notably, the EI signature appeared uncoupled from cognate RNA expression, which instead largely explained differences between monocytes and TAMs.

### SNAI1 activity and cHub associates with fetal-like endothelial cells in PM

Tumor associated stromal cells contribute to cancer growth, metastasis, and resistance to treatment (53), yet their epigenetic heterogeneity has been underexplored, especially in PM. Our scATAC-seq data contained 4,132 stromal cells that formed five distinct clusters corresponding to CAFs, SMCs, normal mesothelium, VECs, and lymphatic endothelial cells (LECs; **Figure 5A, S7A**). Differential motif activity analysis across stromal subsets predicted several TFs with known roles in the cell type where their activity was enriched (**Figure 5B**, **Supplementary Table 2**), including GATA4, a key TF in mesoderm development (30,54), RUNX1, a known regulator of CAFs in colorectal cancer and fibroblast to myofibroblast differentiation (3,55), and MEF2C, a master regulator of SMCs expression (25). The analysis also suggested associations between TFs and cell type-specific regulatory roles that have not been previously reported, such as TCF3 in LECs (**Figure 5B**). We also detected 6,942 cHubs within the stromal compartment (**Supplementary Table 3**), which had predominantly cell type-specific chromatin accessibility patterns and highlighted the epigenomic similarity between mesothelial cells and CAFs (**Figure S7B**). In agreement, mesothelial cells are known to share a mesodermal developmental lineage with fibroblasts and to give rise to myofibroblasts upon lung injury (56).

**Figure 5.**
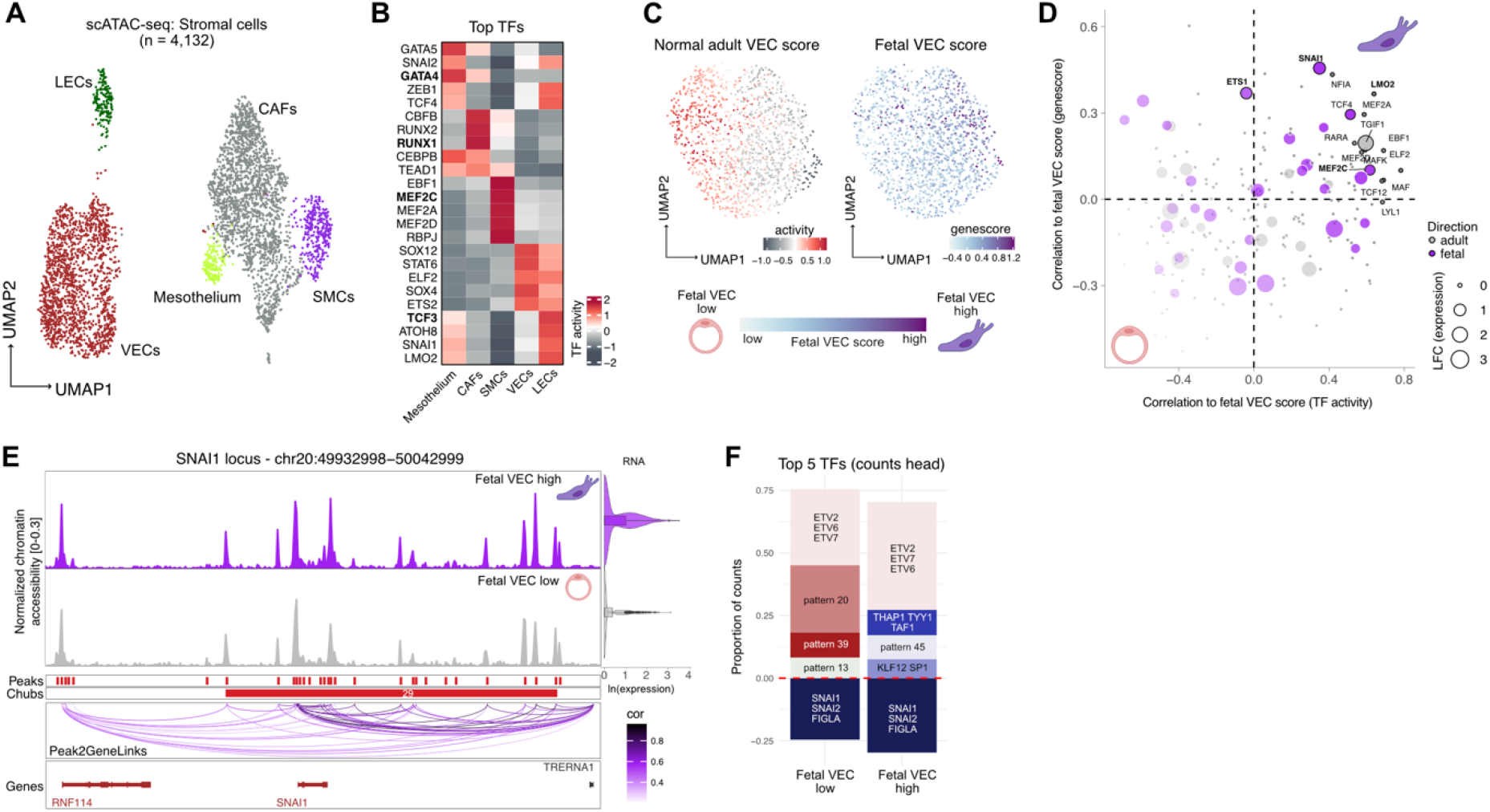
SNAI1 activity and cHub is associated with fetal-like endothelial cells in PM. **A,** UMAP of stromal cell scATAC-seq data colored by cell annotations. **B,** Heatmap displaying the top 5 differentially active TFs in stromal cell subsets. **C,** UMAPs of VECs scATAC-seq data colored by normal adult (left) and fetal (right) VECs score. The fetal VEC score was computed per endothelial cell based on a gene set defined previously (21) (**Methods)**, whereas normal adult VEC score was obtained from (35). **D,** Scatter plot showing the correlation between fetal VECs score and TF genescore (y-axis) and between fetal VECs score and TF activity (x-axis) for all TFs expressed in VECs. **E,** Genome browser at the SNAI1 locus showing pseudobulk chromatin accessibility for cells with high and low fetal VECs scores. Also displayed are SNAI1’s RNA expression (violin plots, right), peak to gene correlation (purple arches, bottom), and cHubs detected (red bars). The number assigned to each cHub indicates its rank by region size with 1 denoting the largest, where cHubs were recomputed within stromal cells. **F,** Top 5 most frequent motifs calls detected from chromBPNet counts outputs after training chromBPNet models using cells with high and low fetal VECs scores. Motifs with negative proportions are associated with negative contributions to chromatin accessibility.

We previously uncovered a fetal-like VEC subset that likely plays a key role in PM angiogenesis (21). To further investigate its epigenomic underpinnings, we first computed a per cell fetal VEC score using a marker gene list we identified previously (21), which highlighted the fetal-like VEC subset in scATAC-seq data (**Figure 5C**, **right**). In agreement, the fetal VEC score was anti-correlated with the epigenome of normal lung adult VECs (35) across PM VECs (**Figure 5C**, left) and across metacells in all PM samples with at least 100 VECs (**Figure S7C, Methods**). Next, to identify the key TF regulators of fetal-like VECs using our single-cell epigenomic data, we correlated the fetal VEC score with TF motif activities and relative genescores (**Figure 5D**). Among the top correlated TFs, we identified SNAI1, which also showed high RNA expression in fetal-like VECs from matched scRNA-seq data and harbored a differentially accessible cHub compared to adult VECs (**Supplementary Table 3, Method**) at its gene locus (**Figure 5D-E**). Snai1-conditional mutant embryos were shown to be lethal at 9.5 days of gestation presenting defects in their vascular network (57), which supports SNAI1’s potential role in reshaping PM vasculature. Several other top correlated TFs that may mediate a transition to a fetal-like VEC state included LMO2 and two previously identified TFs, MEF2C and ETS1 (21), where ETS1 also contained two DAcH flanking its promoter region (**Figure 5D, S7D**). We then trained chromBPNet models on cells with high and low fetal VEC scores, which uncovered ETV TF family motifs, particularly ETV2, as highly enriched in both cell groups (58) and supported SNAI1’s potential role in regulating fetal-like VECs (**Figures 5F, S7E).**

In summary, our scATAC-seq atlas defined the chromatin accessibility landscape and TF activities governing stromal cells and characterized the key cHubs and regulators, including SNAI1, of fetal-like VECs in the PM TME.

### Exhausted CD8 T and KLRC1 NK cells share epigenomic regulatory circuits

We then investigated TILs including 14,674 T and NK single-cell epigenomes **(Figure 6A, S8A)**. Using genescores of canonical markers, we annotated four T cell subsets: CD4, regulatory (Tregs), CD8, and exhausted CD8 T cells (CD8 Ex). To further ascertain our epigenomic-based annotation of the CD8 Ex subset, we performed GSEA analysis on differentially accessed peaks between CD8 Ex and CD8 T cells (Ex-DAPs; **Supplementary Table 7**), which confirmed a significant enrichment for exhaustion-related genes (59) **(Figure S8B-C)**. We also identified two NK cell subsets positive for FGFBP2 and KLRC1 genescores, respectively (**Figure 6B**). The KLRC1 NK cluster represents a NK cell subset previously reported to be enriched in tumor tissues (60,61). This subset displayed higher genescore of inhibitory genes and lower genescore of cytotoxic markers relative to the FGFBP2 NK cluster, as observed in NSCLC (61) **(Figures 6B, S8D)**. Finally, we performed *de novo* clustering on 2,878 B cells in our scATAC-seq dataset, which formed five clusters that we annotated into three cell subsets based on established features (**Figure S8E-H**): memory B (clusters C2-3 enriched for CD80, TOX), naïve B (clusters C4-5 enriched for TCL1A, SPRY1, and TF ELF2 activity), and plasma cells (high POU TF family activity) (62,63).

**Figure 6.**
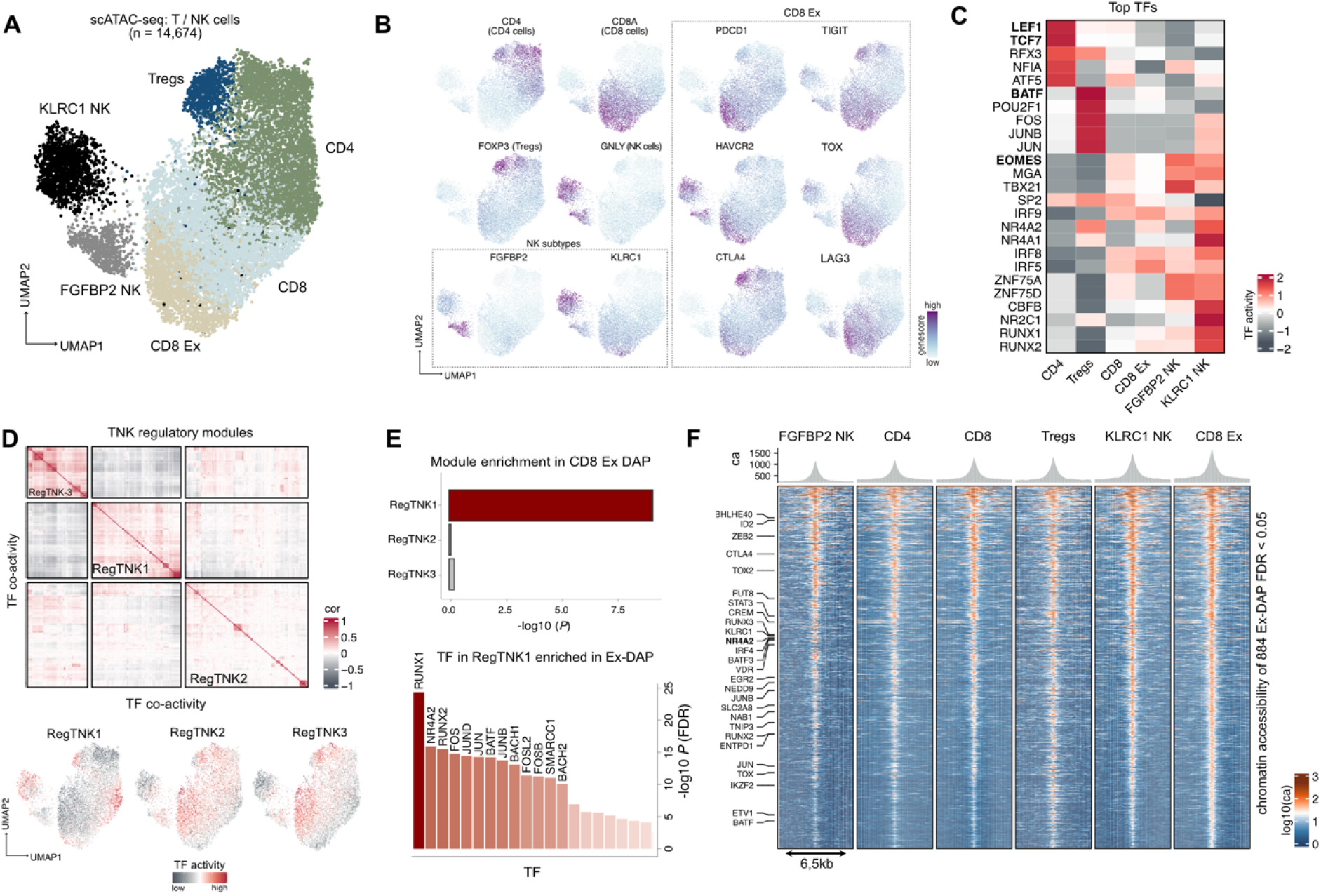
Exhausted CD8 T and KLRC1 NK cells share open chromatin regulatory circuits. **A,** UMAP of T/NK cell scATAC-seq data colored by annotated cell subsets. **B,** UMAPs colored by genescores of established T/NK cell subset markers. **C,** Heatmap displaying the top 5 differentially active TFs across different T/NK cell subsets. **D,** Top: Heatmap of pairwise Spearman correlation between TF activities split into three TF co-activity regulatory clusters (RegTNK1-3). Bottom: UMAP from **A** colored by the average activity of TFs in RegTNK1-3. **E,** Top: Motif enrichment in 884 CD8 Ex-DAP for TFs in RegTNK1. Bottom: TFs in RegTNK1 with motifs most enriched in Ex-DAPs. Enrichment was measured using *P* values from a hypergeometric test (FDR-corrected). **F,** Chromatin accessibility at the 884 Ex-DAPs for each annotated T/NK cell subset. Highlighted are genes of interest located near the peaks or whose gene scores correlate with peak accessibility within 250 kb.

Focusing on T and NK cells, we then performed differential TF activity analysis identifying an enrichment of LEF1 and TCF7 activity in CD4 T cells, BATF motifs in Tregs (64), and EOMES in NK cells (65) **(Figure 6C**, **Supplementary Table 2)**, in agreement with previous scATAC-seq data analyses in basal cell carcinoma (8). Correlation of all TF motif activities across T/NK cells identified three global regulatory clusters **(Figure 6D, top**; **Supplementary Table 5)**. Of these, RegTNK1 consisted of 105 TFs and exhibited the highest average activity in Tregs, a subset of CD4, CD8 Ex, and KLRC1 NK cells **(Figure 6D, bottom)**. We observed significant overlap between Ex-DAPs and peaks harboring RegTNK1 motifs, confirming that cluster RegTNK1 is linked to T cell exhaustion (**Figure 6E**, top; FDR-corrected hypergeometric test), where RUNX1/2, NR4A2, and members of the AP1-complex were among the top enriched TF motifs (**Figure 6E, bottom**). Interestingly, accessibility at the Ex-DAPs was comparable between CD8 Ex and KLRC1 NK cells and less so in the other T and NK cell subsets (**Figure 6F**). This observation remained true when inspecting chromatin accessibility at the top 1,000 DAPs associated with CD8 Ex T cells in four additional human cancers (66) (**Figure S8I**). In sum, scATAC-seq data analysis of TILs suggests that dysfunction-related epigenomic regulatory circuit may be shared between CD8 T cells and KLRC1 NK cells in PM.

### Exhaustion-specific enhancer cRE138020 likely regulates NR4A2 expression

To further investigate the shared epigenomic regulatory circuits of CD8 Ex and KLRC1 NK cells in PM, we sought to identify regulators with enriched activity in both cell subsets. To that end, we ranked TFs in regulatory cluster RegTNK1 based on mean activity in CD8 Ex and KLRC1 NK cells and highlighted those that also exhibited high RNA expression in sample-matched scRNA-seq data, uncovering NR4A2 as a top hit (**Figures 7A**, **S9A**; **Supplementary Table 5**). We also observed higher chromatin accessibility in CD8 Ex and KLRC1 NK cells compared to other T/NK subsets at regions centered on NR4A2 motif instances, providing additional evidence for NR4A2 subset-specific regulatory role (**Figure 7B**). Examining the T/NK-specific *cis-*regulatory landscape, we identified 2,705 cHubs (**Figure 7C**, **Supplementary Table 3**) with several cHubs mapping to the locus of genes with known roles in T cell stimulation and suppression, including PDCD1, HAVCR2, CTLA4, and ICOS (**Figure 7C**). Interestingly, the three cHubs at the CTLA4 and ICOS locus displayed high cell subset specificity of chromatin accessibility patterns, especially for Tregs (**Figure S9B**). At the NR4A2 locus, we identified two cHubs, of which cHub.601 was distally located and harbored cRE138020, a cRE that displayed one of the highest correlations with *NR4A2* RNA expression compared to other cREs within 250kb of the NR4A2 transcription start site (**Figure 7D-E**). The predicted interaction between distal cRE138020 and the NR4A2 promoter was corroborated by independent chromatin conformation capture coupled with immunoprecipitation and sequencing (HiChIP) data from *in vitro* culture of Chimeric Antigen Receptor (CAR)-T cells, which also showed a higher interaction signal in exhausted versus non-exhausted CAR-T cells (**Figure 7D**, **bottom**; **S9C**) (67). Notably, cRE138020 displayed higher accessibility in CD8 Ex and KLRC1 NK cells compared to other T/NK subsets (**Figure 7D**, red arrows) and greater specificity for these subsets when compared to promoter accessibility and NR4A2 expression across all annotated cell subsets in our PM atlas (**Figure 7F**). In addition, cRE138020 had higher accessibility in CD8 Ex versus CD8 T cells and KLRC1 versus FGFBP2 NK cells not only in PM, but also in published NSCLC and clear cell renal cell carcinoma (ccRCC) scATAC-seq data after re-annotation of T and NK cell subsets, suggesting that this predicted regulatory interaction is generalizable among TILs across cancers (10,68) (**Figure 7G, S9D-H**).

**Figure 7.**
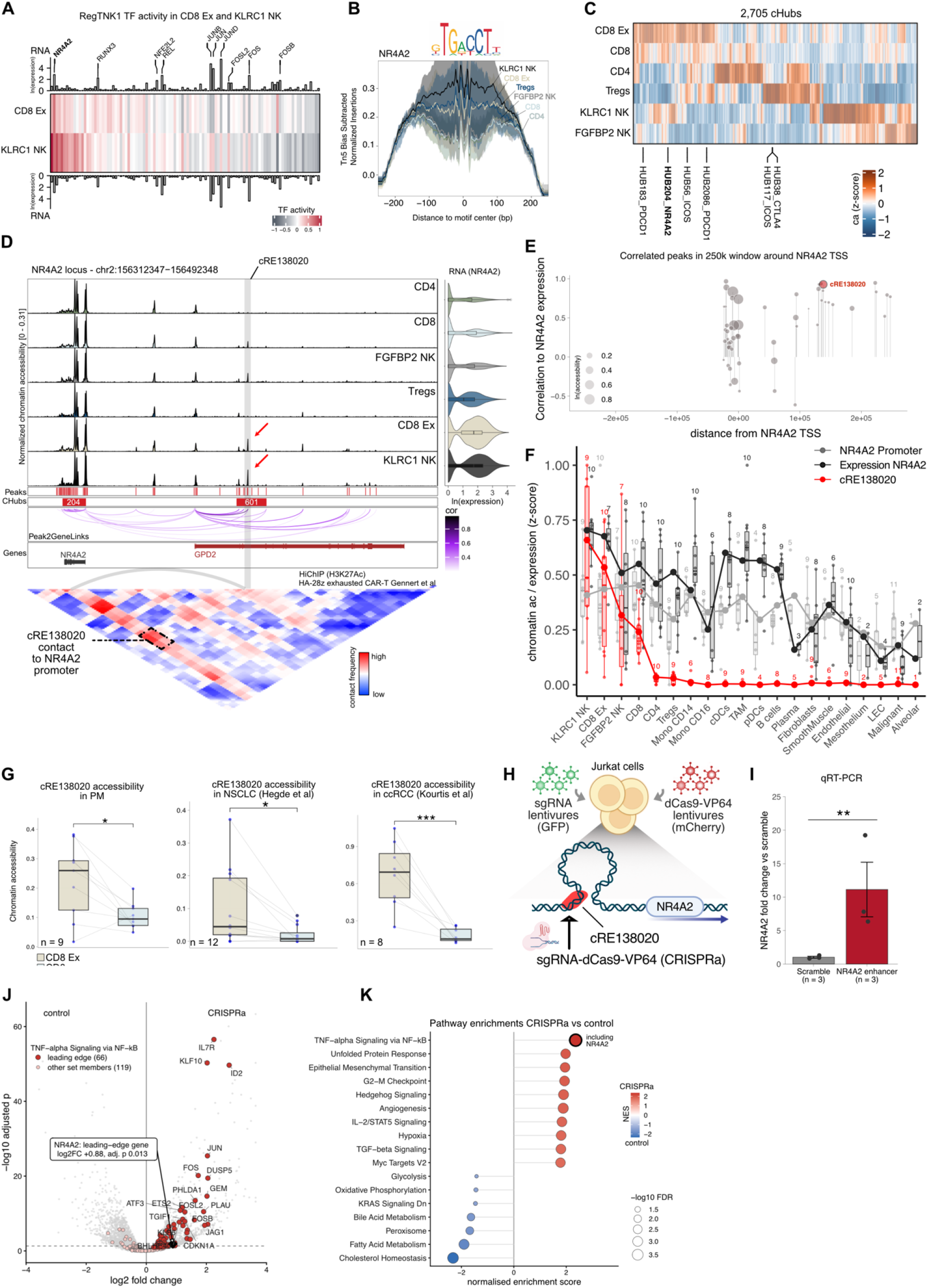
Exhaustion-specific enhancer cRE138020 likely regulates NR4A2. **A,** Heatmap of TFs in RegTNK1 with the highest activity in CD8 Ex and KLRC1 NK. Grey bar plots (top and bottom) show the relative RNA expression of each TF in paired scRNA-seq samples. TFs with highest RNA expression are labelled. **B,** Footprint analysis centered on NR4A2 motif instances (x-axis) showing normalized chromatin accessibility for each T/NK subset (y-axis). **C,** Heatmap of cHubs chromatin accessibility across annotated T/NK cell subsets, where cHubs at loci of interest are highlighted. The number assigned to each cHub indicates its rank by region size with 1 denoting the largest, where cHubs were recomputed within T/NK cells. **D,** Top: Genome browser at the NR4A2 locus showing pseudobulk chromatin accessibility tracks for all annotated T/NK cell subsets. Also displayed are increased accessibility in CD8 Ex and KLRC1 NK subsets at cRE138020 (red arrows), NR4A2’s RNA expression (violin plots, right), peak to gene correlation (purple arches, bottom), and cHubs detected (red bars). Bottom: Interaction heatmap based on Hi-ChIP data from (67) shows contact between cRE138020 region (highlighted in grey) and NR4A2 promoter. **E,** Distribution of peaks in a 250k genomic window centered at the NR4A2 TSS. Height of data points represent correlation between NR4A2 expression and cRE chromatin accessibility computed across T/NK cell subsets scATAC-seq and paired scRNA-seq pseudobulk profiles. Bubble size represents sample mean chromatin accessibility. **F,** Box plots comparing chromatin accessibility at cRE138020 and at NR4A2 promoter regions as well as NR4A2 expression from matched scRNA-seq samples across all annotated cell subsets in the PM scATAC-seq atlas. Also shown are the sample sizes for each track per cell subset. **G,** Sample distributions of chromatin accessibility at cRE138020 compared between CD8 Ex and CD8 T cells in scATAC-seq data from PM (left), NSCLC (middle) (10), and ccRCC (right) (68), where significance was assessed using Student’s *t*-test and n denotes the sample size. **H,** Schematic of *in vitro* validation targeting cRE138020 using CRISPRa in Jurkat cells. **I,** Expression fold-change following qRT-PCR of NR4A2 in cRE138020 CRISPRa-edited Jurkat samples compared to scramble control samples normalized to β-actin expression (n = 3 per condition, Student’s t-test). **J,** Volcano plot showing differentially expressed genes (DEGs) between cRE138020 CRISPRa-edited versus control samples in Jurkat cells. NR4A2 and other genes in its program from panel **K** are labeled. Dashed line represents *P* value = 0.05 on the -log10(FDR adjusted *P* value) y axis. **K,** GSEA pathway analysis on the DEGs identified in panel **J**. Box plot horizontal lines show the 25th, 50th (median) and 75th percentiles, with vertical whiskers extending to a maximum distance of 1.5× the interquartile range from the hinge. Data beyond the whisker ends are plotted individually. *P* values: \**P* < 0.05, \*\**P* < 0.01, \*\*\**P* < 0.001, where *P* values > 0.05 are not shown.

To experimentally test the impact of cRE138020 on NR4A2 expression, we used CRISPR activation (CRISPRa) to target this genomic region using *in vitro* culture of Jurkat cells, an immortalized human T cell line. CRISPRa perturbation demonstrated higher NR4A2 expression when compared to scramble control samples after quantitative reverse transcription polymerase chain reaction (qRT-PCR; **Figure 7H-I, S9I)**. Performing the same experiment with a bulk RNA-seq read-out, we also found significant upregulation of NR4A2 in CRISPRa versus control samples (**Figure 7J**). Moreover, GSEA showed the highest enrichment for HALLMARK gene program “TNF-alpha Signaling via NF-kB” containing NR4A2 (**Figure 7K)**, demonstrating that CRISPR activation of cRE138020 was sufficient to elicit NR4A2 transcriptional activity.

Overall, our high-resolution epigenomic profiling of lymphocyte subsets in PM uncovered previously unappreciated similarities in chromatin accessibility landscapes between CD8 Ex and KLRC1 NK cells and a potential dual role for NR4A2 in their regulation. Intriguingly, distal cRE138020 emerged as a modulator of NR4A2 expression making it an attractive epigenomic target to potentially fine-tune NR4A2 expression.

## DISCUSSION

In this study, we built the first scATAC-seq atlas of the PM TME, *de novo* annotating 24 cell subsets and comprehensively mapping their chromatin accessibility and regulatory landscapes. The high-resolution data allowed us to identify 68 TFs with higher motif activity in malignant versus normal mesothelial cells, where we functionally tested the impact of a subset of these TFs and confirmed overall congruent phenotypes across multiple PM cell lines. We found that genetic perturbation of multiple TFs significantly reduced cancer cell proliferation and all TF KOs displayed dysregulation of distinct cancer-intrinsic pathways (e.g., cell migration) compared to control cells, underscoring the high predictive value of our scATAC-seq data to *de novo* identify putative regulators of PM.

Sarcomatoid tumors are more aggressive and less responsive to most chemotherapies compared to epithelioid PM (42); hence, characterization of their regulatory landscape can aid the design of novel therapies tailored to this more refractory subtype. Our analyses indicate that a putative feed-forward regulatory circuit, including SOX9, RUNX2, and SNAI2, may drive a sarcomatoid-like phenotype in PM. Consistent with our observations, prior work showed that sustained induction of Sox9 transgene in murine epidermal stem cells led to basal cell carcinoma tumorigenesis and activation of RUNX and SNAI family TFs (43). SOX9 oncogenic role has been reported in multiple cancers, including prostate, gastric, and triple-negative breast cancer (69–71), and our work further implicates SOX9 in potentially contributing to a sarcomatoid-like phenotype in PM tumors.

Our data suggest a putative epigenomic-based regulatory dichotomy separating monocytes and MDMs from TRMs. This axis of variation, represented by the EI signature, is not detected by transcriptomics alone, and is associated with AP-1 complex and inflammation-related TFs, such as NF-κB and NFE2L2, with higher motif activities for these TFs in monocytes and MDMs compared to TRMs. We previously showed a similar epigenomic signature in TAMs and bone marrow-derived monocytes in NSCLC and implicated NFE2L2 as a critical player in establishing tumor-driven myelopoiesis (10). Our data further support these findings and highlights that the putative MDM-TRM regulatory axis is uncoupled from cognate RNA expression of associated regulatory programs. Future experiments pairing chromatin accessibility assays with lineage tracing methods in animal models will be required to establish a clear link between the EI signature and macrophage ontogeny.

We have previously shown that NK cells are abundant in PM compared to other cancer types and are associated with better overall survival in patients with epithelioid tumors (21). Here, we further characterized the epigenomes of KLRC1 NK cells, also known as ENTPD1+ tumor-associated NK (taNK) cells, which have demonstrated therapeutic potential upon NKG2A blockade in ex-vivo 3D models of NSCLC (61). Our chromatin accessibility atlas suggests epigenomic similarities between exhausted CD8 T cells and KLRC1 NK cells, extending previously reported parallels between these developmentally related cytotoxic lymphocyte subsets in epigenomic memory acquisition (72), metabolism (73), and at the transcriptional level (74). These findings further support a rationale for functional studies to test whether shared regulatory mechanisms could be targeted to restore cytotoxicity in both T and NK subsets.

NR4A2 is a well-known modulator of key genes affecting CD8 T cell exhaustion and has been proposed as a target to improve fitness of CAR-T cells in *ex vivo* settings (75). Our study uncovered a state-specific enhancer, cRE138020, that is more accessible in exhausted CD8 T cells and KLRC1 NK cells in PM and two other cancer types. Moreover, we demonstrated that CRISPRa of cRE138020 increased expression of NR4A2 and its associated expression program in Jurkat cells. While further experimental evidence in *in vitro* and *in vivo* preclinical models are required, we speculate that regulatory regions such as cRE138020 could be evaluated as targets for improving current cell therapies or *in vivo* approaches to restore endogenous anti-tumor immunity (76). Indeed, targeting cREs rather than their associated genes or proteins may enable more selective and physiological modulation of gene expression, as exemplified by targeting an enhancer controlling PD1 expression, which was reported to avoid the adverse effects associated with complete PD1 knockout in *in vivo* models (12).

Our study has several limitations. First, the CROP-seq and bulk RNA-seq KOs were generated in mesothelioma cell lines, which may not fully recapitulate the cellular programs of malignant cells within the TME. Further validation in mouse models will be needed to ascertain the *in vivo* relevance of these findings. Second, while we demonstrated a link between cRE138020 and NR4A2 expression, additional experiments will be required to functionally test its ability to restore or preserve T and NK cell cytotoxic potential. Finally, we acknowledge the limited sample size of our study, due to the rarity of PM cases. While we obtained enough cells to confidently annotate 24 cell subsets, future cross-institutional studies will likely provide better sampling of less frequent TME cell subsets.

Overall, our scATAC-seq atlas of PM provides insight into the regulatory programs associated with cellular heterogeneity within the TME and a framework for identifying potential therapeutic targets in this rare and lethal disease with limited treatment options. Our findings highlight the epigenome as an important regulatory layer for understanding cellular states and nominate TFs and regulatory elements that could be explored as potential targets for future strategies aimed at modulating the TME.

## METHODS

### Human tumor sample collection

Tumor samples were obtained from diagnostic biopsies and surgical specimens of treatment naïve patients with PM undergoing treatment at Mount Sinai Hospital after obtaining informed consent in accordance with a protocol reviewed and approved by the Institutional Review Board (IRB) at the Icahn School of Medicine at Mount Sinai (IRB Human Subjects Electronic Research Applications STUDY-23-01457) and in collaboration with the Biorepository and Department of Pathology. De-identified clinical information of subjects can be found in Supplementary Table 1.

### Tumor sample processing

Tumor samples were transported in MACS Tissue Storage Solution stored at 4°C, rinsed with PBS, minced, and incubated in a rotation shaker for 40 minutes at 37°C in Collagenase IV 0.25mg/ml, Collagenase D 200U/ml and DNAse I 0.1mg/ml (all Sigma). Cell suspensions were aspirated through a 18G needle ten times and strained through a 70-micron mesh prior to red blood cell lysis. Dead cells were removed using an EasySep Dead Cell Removal (Invitrogen, V13242) Kit. For surgical resections, cell suspensions were additionally sorted into CD45+ and CD45- cells using the EasySep™ Human CD45 Depletion Kit (StemCell, 17898) per kit instructions. Next, CD45+ and CD45- sorted cell populations were pooled in equal numbers to ensure sufficient sampling of all cell types in the PM TME; however, this cannot ensure even numbers for each cell class, since cell viability can vary between cell types and will ultimately affect the number and identity of sequenced cells. For biopsy samples, we did not carry out CD45 sorting due to insufficient starting cellular material.

### Tumor single-cell library construction and sequencing

For scATAC-seq profiling, cells were washed and resuspended in nuclei lysis buffer (10 mM Tris-HCL (pH 7.4), 10 mM NaCl, 3 mM MgCl2, 0.1% Tween-20, 0.1% nonidet P40 substitute, 0.01% digitonin, 1% BSA) and incubated on ice for 4-5 min, followed by addition of chilled wash buffer (10 mM Tris-HCL (pH 7.4), 10 mM NaCl, 3 mM MgCl2, 1% BSA, 0.1% Tween-20). Nuclei were pelleted by centrifugation, resuspended in 1X nucleus buffer (10x Genomics, 1000493) and counted using Acridine Orange/Propidium Iodide viability staining reagent (Fisher Scientific, NC1765655) and Cellometer Auto2000 Automated Cell Counter (Nexcelom). As expected, following lysis, the AO/PI viability readout was <1%, consistent with efficient nuclei release. ScATAC seq was performed using 10x Genomics’ Chromium Next GEM Single Cell ATAC Reagent kit v1.1, following manufacturer’s instructions. Libraries were amplified by PCR, and final library size distribution and concentration were assessed using an Agilent Technologies Bioanalyzer and Thermo Fisher Scientific Qubit, respectively. Libraries were then sequenced on an Illumina HiSeq 2500 or Illumina NovaSeq instrument. ScRNA-seq profiling of one additional sample (P14) was performed as described previously for other scRNA-seq samples in our cohort (21).

### ScATAC-seq data processing and analysis

Pair-ended FASTQ files were aligned to hg38 genome reference using CellRanger software (cellRanger-atac v1.2.0). Raw peak matrices and fragment files were initially input in Signac (v1.14.0) (77) for QC and cell type annotation for each individual sample. Cells that passed all of the following QC metrics were retained for further analysis: > 3,000 peaks, < 30,000 peaks, > 15% of reads in peaks, < 0.05 ratio of blacklisted peaks, < 4 nucleosome signal, and >3 enrichment at transcription start sites (TSSs). Data normalization and linear dimensionality reduction was performed as per standard pipelines using term frequency–inverse document frequency (TF-IDF) normalization followed by singular value decomposition (SVD). Non-linear dimensionality reduction was performed using uniform manifold approximation and projection (UMAP) followed by construction of a nearest-neighbors graph.

To cluster cells, we used Louvain community-detection algorithm (78) as implemented in Seurat FindClusters (algorithm 1, standard modularity), called through ArchR addClusters for scATAC-seq and directly for scRNA-seq. Briefly, a k-nearest-neighbor graph was built on the reduced dimensions (latent semantic indexing (LSI) or Harmony-corrected LSI for scATAC-seq, k = 20; first 15 principal components for scRNA-seq, k = 30) and converted to a shared-nearest-neighbor graph weighted by the Jaccard overlap of neighborhoods, pruning edges below 1/15. Modularity was optimized from multiple random starts (10 for scATAC-seq, 100 for scRNA-seq; 10 iterations each), keeping the partition with the highest modularity. Resolution was set per dataset (0.6–3 for scATAC-seq; 0.8–1.2 for scRNA-seq). For scATAC-seq, clusters with fewer than 5 cells were reassigned to the cluster of their 10 nearest neighbors. Cell type annotation was guided by transferring labels from pre-annotated sample-matched scRNA-seq (label transfer) (79), and examination of the genescore and TF activity of *bona fide* cell type markers and TFs, respectively.

For other scATAC-seq analyses, we used ArchR (v1.0.2) (80) as it provided a computationally more efficient alternative to Signac. Fragment files for each sample were used as input and then filtered by the high-quality cells previously identified with Signac. Combined data was similarly processed in ArchR using *addIterativeLSI* to reduce dimensionality and *addUMAP* generate UMAP graphs. Cellular compartment UMAPs were batch-corrected using harmony (81). Peaks were called using MACS2 (v2.2.9.1) via *addReproduciblePeakSet* function and were retained only if detected in more than one pseudo-bulk group coverage except for peaks called in malignant clusters. TF activity was measured using chromVAR (v1.24.0) (22) using CIS-BP motif database (82). TFs with ln(RNA expression) < 0.1 in the corresponding cell type were considered lowly expressed and filtered out from downstream analyses. Footprint analysis was performed using *getFootprints* and *plotFootprints* with Tn5-bias subtraction. Differential TF activity, peak accessibility and genescore was performed using ArchR function *getMarkerFeatures*. To compute gene module scores from gene sets of interest the function *addModuleScore* was used.

GSEA was performed with fgsea (v1.28.0) on gene-score differences from getMarkerFeatures (Wilcoxon test, bias-matched background cells). Genes were ranked by log2 fold change and tested against MSigDB GO Biological Process gene sets (fgseaMultilevel; minimum set size 15). Significant pathways (adjusted p < 0.05) were collapsed to non-redundant sets with collapsePathways.

### Sc-ATAC-seq analyses of PM malignant cells

To detect TF motifs enriched in tumor cells compared to normal mesothelial cells we first combined PM malignant cells with normal mesothelial cells from distal normal lung (35) in one ArchR project. For each TF motif, chromVAR deviations of each tumor’s cells were compared with those of the normal mesothelial cells by a Wilcoxon rank-sum test with FDR correction. Motifs were retained if they were significantly higher in tumor cells (FDR < 0.05) in at least 5 of the 11 tumors. TFs were further required to be expressed (log2(mean expression + 1) > 0.5) in scRNA-seq of at least one of 9 tumors (P10 did not have any malignant cells in scRNA-seq) and higher mean RNA expression in tumor cells than in normal pleural mesothelium. This yielded 68 TF motifs reported in **Supplementary Table 4**. For scATAC-seq analyses performed on metacells of malignant clusters, we excluded samples P3 and P13 due to insufficient cell numbers.

### Correlative analysis of epigenomic features

For TF co-activity analysis, activity of each TF pair was correlated across cells using Spearman correlation and TF modules were identified using K-means clustering with R function *kmeans.* For associating TF activities to gene module scores (e.g., TF activity vs scS-score), Spearman correlation was computed per sample between the two variables examined across metacells (defined as the average point of a group of cells sharing similar features) for that sample. Metacells were generated by binning barcodes pre-sorted by the module score of interest and averaged using function *rollapply* from package zoo (v1.8.12) with bins including 30 to 50 cells depending on the overall number of cells. Module scores were computed using ArchR addModuleScore of the gene sets using genescores. For finding association between pairs of TF activities Spearman correlations were computed directly across single cells. The median value of the Spearman correlations computed in each sample was used to rank most relevant associations. The scS-score was derived using the sample cohort and method in (21) but recomputed to include malignant cells from additional sample P14.

VEC cells were scored with VEC subsets module scores identified previously (21) including a fetal-like (COL4A1□), arterial and venous subsets. Module scores were z-scored across the three modules within each cell, and cells were split at the per-sample median of the fetal VEC score into high and low groups. The EI signature was defined as the average of top 14 TF activities inversely associated with PC1 (**Figure 4D)** and with activity correlated with relative genescore (i.e., activating TFs). Myeloid cells were either labeled EI score high (EI score > 0) and low (EI score ≤ 0), where EI and other gene sets are reported in **Supplementary Table 5**.

### Chromatin hubs detection algorithm

To detect cHubs, we adapted the Cicero co-accessibility algorithm (31) to identify clusters of co-accessed peaks. To do so, we first computed peak co-accessibility on metacells generated with a customized version of ArchR function *addCoaccessibility* and then stitched together all adjacent co-accessed peaks with distance ≤ 12.5 kb. This parameter was previously set to detect SE regions from ChIP-seq data using the ROSE algorithm (28). In addition, co-accessed peak pairs were filtered based on Pearson correlation (*r* > 0.2 - 0.3). The resulting co-accessibility matrix was then converted in a graph using igraph package (v2.0.3) and cHubs were defined as genomic regions having ≥ 5 co-accessed peaks. This minimum peak count was chosen to prioritize detection of larger regulatory regions. For downstream analyses using cHubs, such as DAcH, the fragments sum spanning the whole cHub region normalized by number of reads in TSS (ReadsInTSS) was used. We run our cHub detection algorithm across the consensus peak set spanning all 49,849 cells. When analyzing specific cell compartments in the manuscript, we also ran the algorithm on the corresponding cell type-specific peak sets to improve detection of cell subset cHubs, where algorithm run iterations with relative parameters settings are reported in **Supplementary Table S3**.

### Peak and cHubs gene annotation

To map peaks and cHubs to genes they may regulate we used their nearest gene from the ArchR peak annotation (distance to gene start). For mapping Ex-DAPs to genes they may regulate, we further used addPeak2GeneLinks on the T/NK cell compartment. This correlates peak accessibility with gene scores across metacells in the IterativeLSI space, for peak–gene pairs within 250 kb. Links with correlation ≥ 0.35 and FDR ≤ 1e-4 were retained, and each peak was assigned the gene with its strongest link. Peaks without a retained link (207 of 884) were assigned their nearest gene from the ArchR peak annotation.

### Overlap of cHubs and SE regions and benchmarking

To compute overlap of detected cHubs regions from our scATAC-seq dataset with SE regions obtained from the SE.db database (https://bio.liclab.net/sedb/, accessed 09/01/2023), we utilized a hypergeometric test with *phyper* R base function, where *q* represents the intersection of SE regions with cHubs, *m* the intersection of SE regions with background peaks, *n* the (number of background peaks) – *m*, and *k* the number of cHubs identified. The test was repeated for cHubs identified from each PM cell type in our scATAC-seq dataset with SE regions from each primary cell type in the SE.db database. Background peaks corresponded to all peaks detected for a given cell type. *P* values were FDR-corrected.

We benchmarked our cHubs detection method and its robustness to different parameter settings by directly comparing it to stitching proximal peaks together after calling cell type-specific peaks (Stitch-CT) or starting from the consensus peak set (Stitch-AP). For the Stitch-AP comparison, per cell type peak hubs were further identified via DAcH. Both methods were run across peak distances of 5, 10, 12.5, 15 and 20 kb, minimum hub sizes of 3, 4 or 5 peaks, and with or without excluding peaks within 2 kb of a TSS. Overlap of these hubs with SE sets (SE.db) was tested against the cell type’s reproducible peaks with a one-sided hypergeometric test and FDR adjusted within each setting. Eight pairs of cell type and matching SE set were used for assessing the performance of each algorithm run: alveolar with AT1 and with AT2, CAFs with lung fibroblast, SMCs, VEC with endothelial, myeloid with CD14 blood monocyte, T cells, and B cells. SE recovery was defined as the mean −log10 FDR of the matched pairs, each value capped at the 99th percentile and set to 0 for a pair without differentially accessible hubs. SE specificity was defined as recovery minus the mean −log10 FDR of the same cell types against non-matching SE sets.

### ScRNA-seq data analysis

The pre-processed Seurat object of scRNA-seq data from matching samples (n = 9) was obtained from GSE190597. Raw count matrices from the newly generated scRNA-seq sample (P14) were imported in R using Seurat (v5.1.0) and low-quality cells with unique molecular identifiers (UMIs) < 1,000, gene detected < 400, mitochondrial index > 25% were removed and data was normalized. Sample P14, collected and sequenced as part of this study, was integrated with the rest of the scRNA-seq cohort and annotated cells as done previously (21). We used scRNA-seq data throughout the paper to filter out lowly expressed TFs and as supportive evidence for our scATAC-seq-derived predictions. To identify cancer-related molecular pathways that a given TF might regulate, we generated metacells of malignant cells using hdWGCNA (v0.2.18) (83) and then performed a pathway enrichment test using clusterProfiler (v 4.10.0) function *enricher* and HALLMARK annotation (GSEA-MSigDB v7.4) of the top 100 correlated genes for each TF. Metacells, gene correlation, and enrichment tests were performed per sample to ensure robustness of associations.

For TAM subsets identification in scATAC-seq data, we first ran cNMF (v1.3.4) on the scRNA-seq cohort per sample within the myeloid compartment only. We selected K = 10 cNMF modules for each sample iteration and then grouped all 100 cNMF modules based on their gene set overlaps (using only top 50 genes in each module) in 10 k-means meta-modules (k = 10). Meta-modules that included less than 3 cNMF modules were discarded in so retaining only robust myeloid meta-modules. To investigate gene regulation from scRNA-seq, we inferred transcription-factor regulons in myeloid cell scRNA-seq data using pySCENIC (84) with standard settings on raw UMI counts of 5,000 most variable genes.

### Copy number alteration inference and subclonal analysis

To infer CNA from scATAC-seq we binned the genome of single cells in 10Mbs windows with a sliding window of 5Mbs and compute normalized fragment counts in each bin with a customized version of functions *makeWindows* and *scCNA* from (https://github.com/GreenleafLab/10x-scATAC-2019). Z-scores of the fragment coverage per bin was further averaged per sample. For inference of CNA from scRNA-seq we used inferCNV R package (v1.20.0) (85). For inferring CNA from bulk RNA-seq of mesothelioma cell lines, expression data was binned into 10 Mb windows and for each sample, expression (log2) was averaged within each window and the window values were z-scored across the genome. This gives a genome-wide dosage profile in which gained regions read as relatively high and lost regions as relatively low. The same profile was built from CCLE expression data for NCI-H2052, MSTO-211H, NCI-H2452 and NCI-H28. Each sample was assigned to the CCLE line whose profile correlated best with its own (Pearson r).

Copy-number alterations for subclonal analysis were inferred per tumor from malignant-cell scATAC-seq fragments with the epiAneufinder() function (v1.1.5) (27), using windowSize = 5e6, minFrags = 5000, k = 4, gc_correction = TRUE, genome = “BSgenome.Hsapiens.UCSC.hg38”, exclude = c(“chrX”,“chrY”,“chrM”) and the ENCODE hg38 blacklist (v2). Nine tumors with sufficient malignant cells were analyzed using this approach. Subclones were defined by hierarchical clustering of the per cell copy-number calls (dist(), Euclidean; hclust(), method ward.D2) and cutting the tree in two (cutree(k = 2)). The split was accepted only if both groups contained at least 20 cells and 5% of the tumor’s cells and differed by ≥ 0.55 in mean copy-number profile (fraction of cells with a gain minus fraction with a loss) on at least one chromosome arm; otherwise, the tumor was treated as a single clone. TF motif activity differences between subclones were computed for each of the five tumors with two accepted subclones (P4, P5, P10, P11, P23), between cells of the two clones with a two-sided Wilcoxon rank-sum test. *P* values were FDR adjusted over all motifs within each tumor. In tumors with matched scRNA-seq data, a motif was tested only if its TF was expressed (log2 (mean expression + 1) ≥ 0.5) in the tumor’s malignant cells.

### Visium spatial transcriptomics data collection, processing, and analysis

Four PM tumor tissue sections—two fresh frozen (A and B) and two snap-frozen (C and D)—from tumor P4 were profiled using 10x Visium spatial transcriptomics technology (Spatial 3′ v1 chemistry). Reads were aligned to GRCh38 (10x reference 3.0.0) and transcripts were assigned to spots with Space Ranger 1.0.0. Each section was normalized with SCTransform (Seurat; 3,000 variable features), followed by PCA (50 components) and UMAP and graph-based clustering. Spot cell type composition was estimated using Seurat label transfer method, where P4 patient-matched scRNA-seq data annotated uniformly with other samples in the study was used as a reference. To infer malignant clonal spatial segregation, malignant spots (spots with highest malignant cells prediction scores from label transfer) were assigned a chr8q score computed as the mean log-normalized expression of genes on chr8q minus that of all other autosomal genes, centered on the non-malignant spots of the same section. Malignant spots were split into chr8q-high and chr8q-low groups by k-means (k = 2) and spatial colocalization was tested by comparing neighbor-label agreement with 1,000 label shuffles.

### Acquisition and analysis of bulk RNA-seq data from independent human PM cohorts

RSEM-normalized count matrix including 82 bulk RNA-seq samples as part of the Hmeljak et al. cohort was downloaded using the R package cgdsr (v1.3) of cBioPortal (http://www.cbioportal.org), an online database built for cancer genomics along with metadata including histology information. The Bueno cohort was downloaded from the European Genome-phenome Archive (EGA) under accession number EGAS00001001563 as RPKM-normalized count matrix, including 216 bulk RNA-seq samples along with metadata information including histological and molecular subtypes. TPM-normalized bulk RNA-seq from the Mangiante et al. cohort was obtained from https://github.com/IARCbioinfo/MESOMICS_data (accessed on 04/27/2023). Datasets were log2-normalized before downstream analysis. The association of SOX9, RUNX2 and SNAI2 expression with overall survival was tested in the three bulk RNA-seq cohorts with survival time expressed in months. Because sarcomatoid PM subtype is itself prognostic, each model was fitted both with no adjustment and adjusted for the scS-score. Cox proportional-hazards models were fitted with standardized expression as a continuous variable (hazard ratio per standard deviation) and with expression split at the cohort median (high versus low). For the median split, unadjusted Kaplan–Meier curves were compared with the log-rank test. ScS-adjusted survival curves were obtained by direct standardization from the Cox model, with 95% confidence intervals from 500 bootstrap resamples.

### ChromBPNet models training

To infer TFs associated with counts and distribution of fragments in peaks, we trained chromBPNet models for each cell type of interest (34). ChromBPNet generates two complementary outputs: one predicting the total counts in peaks (count head) and a second one predicting the shape of the count distribution over peaks (profile head). ChromBPNet models were trained on 5 different folds of set of chromosomes split into training and validation sets. Tn5-bias models were trained on T/NK cells and used to regress out Tn5-associated bias for all other cell type models. QC reports for each training round were examined to ensure no residual Tn5-bias-related motifs were significantly enriched in the models. Contribution scores from counts and profile heads were then each averaged across training fold and passed to TFmodisco-lite (v1.0.0) for identification of seqlets and their matching to known TF motifs (HOCOMOCO v11 TF motif database, accessed on: 05/01/2025). Similarly, bigwig files from each fold and head were averaged using wiggletools (v1.2.11) *mean* function followed by *wigToBigWig* function. Finally, we used Fi-NeMo (v0.25) to call all instances of identified motifs for each output head. To visualize contribution score bigwig files the WashU Epigenome Browser was used (https://epigenomegateway.wustl.edu/).

### Reference mapping of myeloid cell subsets

Myeloid cell subsets were annotated by reference mapping onto the scATAC-seq hematopoiesis atlas of (52) (GSE139369), which comprises healthy bone marrow mononuclear cells, CD34 - enriched bone marrow, peripheral blood mononuclear cells and mixed-phenotype acute leukemia samples. A reference ArchR project was built from the published fragment files and dimensionality-reduced by LSI. The authors’ BioClassification annotation (26 hematopoietic clusters plus six mixed-phenotype acute leukemia (MPAL) classes) was used as the label set. All 6,394 cells of the mesothelioma myeloid compartment were projected into the reference LSI space in a single run. For each query cell, the k = 25 nearest reference cells were identified by Euclidean distance in that space. Each cell was assigned the majority label among its 25 neighbors, and a soft score was computed for every reference label as the fraction of the 25 neighbors carrying it, giving values in [0, 1] that sum to one per cell. The score of the assigned label was retained as a transfer confidence, and the mean distance to the 25 neighbors was retained as a mapping-quality measure.

### Hi-ChIP processing and analysis

FASTQ files were downloaded from Sequence Read Archive (SRA) data (accession no. SRP310567) (67) aligned to hg38 genome reference and processed using Hi-C Pro pipeline (v3.1.0). Raw contact maps were normalized using *normPerExpected* function with method ‘loess’, binned in 10kb windows using function *binningC,* replica samples were averaged, and region of interest was visualized with *mapC* function using R packages HICExperiment (v0.99.9) and HiTC (v1.50.0).

### DepMap data acquisition

Bulk RNA-seq and CNA data of mesothelioma cell lines was obtained from https://depmap.org/portal.

### Cell Culture Conditions

Human pleural mesothelioma cell lines NCI-H2452 (ATCC, CRL-5946), NCI-H2052 (ATCC CRL-5915), MSTO-211H (ATCC, CRL-2081) and NCI-H28 (ATCC, CRL-5820) were obtained from the American Type Culture Collection (ATCC) and maintained at 37°C with 5% CO in ATCC-formulated RPMI 1640 (ATCC, 30-2001), supplemented with 10% fetal bovine serum (Gibco, 16000044), and 100 U/mL penicillin–streptomycin (Gibco, 15140148). HEK293T cells (ATCC, CRL-3216) were maintained at 37°C with 5% CO in Dulbecco’s Modified Eagle Medium (Gibco, 11965092), supplemented with 10% fetal bovine serum (Gibco, 16000044), and 100U/mL penicillin–streptomycin (Gibco, 15140148). Antibiotics were omitted for at least 24 hours prior to transfection. Jurkat cells, clone E6-1, (ATCC, TIB-152) were obtained from the ATCC. Jurkat cells were maintained at 37°C with 5% CO in RPMI 1640 (Corning, 10-104-CV), supplemented with 10% fetal bovine serum (Gibco, 16000044), 1x minimum essential medium (MEM) nonessential amino acids solution (Gibco, 11130051), 1mM sodium pyruvate (Corning 25-000-CI), 10mM HEPES (Cytiva, SH40003.01) and 100U/mL penicillin–streptomycin (Gibco, 15140148). Jurkat cells were maintained at a density of 1 million cells/mL. All cells were tested upon receipt and every 2-3 passages for mycoplasma using Plasmotest Mycoplasma Detection kit according to manufacturer’s instructions (Invivogen, rep-pt1) and were negative for contamination.

### Generation of Monoclonal Cas9-Expressing Pleural Mesothelioma Cell Lines

Lentivirus was generated by transfecting 293T cells with lentiCas9-Blast (Addgene, 52962), psPAX2 (Addgene, 12260), and pMD2.G (Addgene, 12259) at 2:2:1 ratio using Lipofectamine 3000 and Opti-MEM I (Gibco, 11058021) according to the manufacturer’s instructions. At 24 hours post-transfection, medium was exchanged for a 1:1 mixture of complete RPMI 1640 and DMEM without antibiotics. Viral supernatant was collected 48 and 72 hours post transfection and pooled prior to being centrifuged (2100 RCF for 5 minutes) and filtered (0.45μm) and immediately added to the PM cell lines for 48 hours. PM cells were selected in cell culture medium supplemented with blasticidin at a final concentration of 5μg/mL for a total of 10 days with regular media changes. Blasticidin-resistant colonies were expanded for approximately 2 weeks in cell culture medium with 2μg/mL blasticidin after which they underwent sorting by a FACSymphony S6 Cell Sorter (BD Biosciences) to be plated as single cells in a 96-well tissue-culture plate. Plates were manually inspected to ensure one event per well after which clones were expanded. Following antibiotic selection, cells were maintained in complete RPMI medium with 1μg/ml blasticidin to maintain selective pressure.

### Verification of Cas9 Expression via qRT-PCR

Once clones had expanded to sufficient cell numbers, total RNA was individually isolated from several clones using the RNeasy mini kit and following the provided protocol (Qiagen, 74104) and the resulting RNA was reverse transcribed with the Lunascript RT Supermix Kit (NEB, E3010L) according to the respective manufacturer’s instructions. QRT–PCR was performed to assess Cas9 mRNA abundance. Reactions were carried out using SYBR Green chemistry in a total volume of 10µL, comprising 5µL SYBR Green Master Mix (Applied Biosystems, 4309155), 1µL 100nM forward primer, 1µL 100nM reverse primer, 1µL cDNA template (diluted 1:3 v/v with nuclease-free water), and 2µL nuclease-free water in triplicate. Thermal cycling was performed on a QuantStudio 3 Flex Real-Time PCR System (Applied Biosystems) with an initial denaturation at 95°C for 10 min, followed by 40 cycles of denaturation at 95°C for 10s and extension at 60°C for 60s. Two previously validated primer sets were used to quantify Cas9 mRNA expression, and expression levels were normalized to β-actin using the ΔΔCt method. Cells derived from the clone exhibiting the highest Cas9 mRNA expression were selected for all subsequent experiments. All primer sequences are found in **Supplementary Table 8**.

### CROP-seq Guide Vector Preparation

Vector backbone preparation mix consisted of 5μg CROPseq-Guide Puro plasmid (Addgene, #86708), 2μL BsmBI (NEB, R00739S) in 2.5μL NEBuffer 3.1 (NEB, B6003S) with nuclease-free water up to a final reaction volume of 25μL. The plasmid was then digested for 1 hour at 55, followed by 20 minutes at 80 to inactivate the restriction enzyme. The resulting digestion was run on a 0.8% agarose gel with SYBR Safe DNA gel stain (Invitrogen, S33102) for 40minutes at 100V, and the 8,333bp fragment was extracted and purified with the QIAquick Gel Extraction Kit (Qiagen, 28704) according to the manufacturer’s instructions.

### gRNA design and Gibson Assembly

Three gRNA sequences per target transcription factor were selected from the Brunello CRISPR Knockout Library, on the basis of highest predicted on-target activity (Rule Set 2 scores). Nontargeting control guide sequences also come from this library but do not have a Rule Set 2 score due to lacking an intended genomic target. Oligonucleotides encoding gRNAs were synthesized (Integrated DNA Technologies) with additional homology overhangs for cloning into the CROPseq-Guide-Puro backbone following this sequence:

1. a 5′ homology prefix (TGGAAAGGACGAAACACCG), including a guanine immediately upstream of the gRNA sequence
2. the gRNA sequence
3. a 3′ postfix (GTTTTAGAGCTAGAAATAGCAAGTTAAAATAAGGC) homology sequence.

Oligonucleotides were diluted to 100nM in EB buffer and individually assembled into the CROPseq-Guide-Puro backbone using Gibson isothermal assembly for 2 hours at 50 °C, with the following reaction composition: 11fmol backbone, 200fmol ssDNA oligo, 10µL NEBuilder HiFi DNA Assembly Master Mix (NEB, E2621S), and nuclease-free water to 20µL. A 3µL aliquot of the assembly reaction was transformed into DH5α E. coli Mix & Go! Competent Cells (Zymo Research, T3009) and incubated on ice for 5 min, followed by plating on LB agar plates supplemented with 100 µg/mL carbenicillin. Plates were incubated overnight at 30°C, and individual colonies were picked and expanded in LB broth containing 100µg/mL ampicillin for 16–20 hours. Plasmid DNA was isolated using the QIAprep Spin Miniprep Kit (Qiagen, 12123) according to the manufacturer’s instructions, and whole-plasmid sequencing was performed to confirm proper gRNA cassette assembly.

### Lentivirus Generation and Transduction of Cas9-Expressing PM Cell Lines

For each target, lentivirus was generated by transfecting HEK293T cells with pooled gRNA plasmids (three per gene) together with packaging plasmids (pMDLg/pRRE, pRSV-Rev, and pMD2.G) at a molar ratio of 2:1:1:1, using Lipofectamine 3000. Viral supernatant was collected following the incubation periods and harvesting protocol detailed above. Cas9-expressing PM cell lines were seeded 24 hours prior to addition of lentivirus in 6-well tissue culture plates at a density of 600,000 cells per well in complete culture medium without antibiotics. Wells were transduced with harvested lentivirus at 1:3 v/v dilution immediately following collection and filtration. After 48 hours, medium was exchanged for complete RPMI culture medium with penicillin-streptomycin for 48 hours prior to starting antibiotic selection with puromycin. Once selection medium (complete RPMI media with 4μg/ml puromycin) was added, the media was changed every 2-3 days for 7-10 days. Following antibiotic selection, cells were maintained in complete RPMI medium with 1μg/ml puromycin and 1μg/ml blasticidin to maintain selective pressure for successfully transduced Cas9-expressing cells.

### Sample Collection prior to CROP-seq and bulk RNA-seq

Cells were detached with 0.25% Trypsin-EDTA (Gibco, 25200056) and collected by centrifugation at 300 RCF for 5 minutes. Cells were washed once with PBS containing 0.2% BSA, resuspended, filtered through a 40μm cell strainer, and manually counted with a hemocytometer. This process was repeated for cells transduced for each target individually. For single-cell sequencing, equal numbers of NCI-H2052 cells from each condition were then pooled to generate a final suspension of 16,000 cells. For bulk-RNA sequencing, 1 million cells were collected per perturbation for each cell line for total RNA extraction with the Qiagen RNeasy Mini Kit (Qiagen, 74104) according to the manufacturer’s instructions.

### Single-cell CROP sequencing

Single-cell viability was assessed to be 95.6% using Acridine Orange/Propidium Iodide viability staining reagent (AO/PI) (Nexcelom). scRNA-Seq was performed using the Chromium X system (10x Genomics) with the 3’ gene expression (3’ GEX) V3.1 kit, targeting recovery of ∼10,000 cells. Gel-Bead in Emulsions (GEMs) were generated on the Chromium X chip and barcoded cDNA was extracted post-GEM RT-cleanup and amplified for 11 cycles. For GEX libraries, amplified cDNA was fragmented and subjected to end-repair, poly A-tailing, adapter ligation, and 10x-specific sample indexing following the manufacturer’s standard protocol. For gene expression library preparation, amplified cDNA was fragmented, end-repaired, A-tailed, ligated to sequencing adapters and indexed using 10x Genomics sample indices according to the manufacturer’s instructions. The gRNA region—intrinsically encoded in the CROP-seq vector—was amplified by PCR using the NEBNext Q5U kit (NEB, M0597S) with vector-specific primers.

F_CROPseq_PCR1:

/5Biosg/UAUAGTGACTGGAGTTCAGACGTGTGCTCTTCCGATCTCGATTTCTTGGCTTT ATATATCTTGTG

AAO272:

CTACACGACGCTCTTCCGAT*C*T.

A total of 8 cycles of amplification were performed. Following PCR, the amplicons were isolated with MyOne streptavidin beads (ThermoFisher, 65601) and purified with SPRI selection (1X, Beckman Coulter). Final indexing PCR was performed using NEB Q5 PCR kit (NEB, M0492L) with Illumina Universal adapters P5 and P7. A total of 27 cycles were performed. Custom sequencing primer (CGATTTCTTGGCTTTATATATCTTGTGGAAAGGACGAAACACCG) was used for sequencing. Sequencing was performed on an Illumina platform (NovaSeq X for GEX and NextSeq for CROPseq), configuring read pairs to capture cell barcodes, UMIs, and mRNA or gRNA sequences. Raw sequencing data were processed using standard Cellranger pipelines (10x Genomics) for gene expression mapping, and to extract and assign gRNA identities to each cell. Downstream analyses integrated guide assignments with gene expression profiles to derive perturbation-specific transcriptional effects.

### CROP-seq data analysis

Raw count matrices were imported into R using Seurat (v5.1.0). Low-quality cells were removed (UMI < 800, genes detected < 400, mitochondrial fraction > 25%), and data were normalized with *NormalizeData*. Cells were retained if all detected guides targeted the same gene, or a single gene in combination with NTC guides. Guides detected in fewer than 10 cells were removed. Guides targeting the same gene that showed concordant average expression profiles on PCA plot were merged for all downstream analyses. Knockout was assessed by comparing target-gene expression between knockout and NTC cells. Cell-cycle phase scores were computed with Seurat *CellCycleScoring* and compared between each knockout and NTC cells (Wilcoxon rank-sum test, FDR adjusted). Activity of the 20 malignant PM modules (Cm1–Cm20) was scored per cell with Seurat AddModuleScore using the top 50 genes of each module (21). Copy-number profiles were inferred with inferCNV (v1.20.0) (cutoff 0.1, denoising, chrX excluded) using normal mesothelial cells as reference.

### Generation of Monoclonal Dual Transduced Jurkat Cells

pZR112_Lenti-SFFV-mCherry-2A-dCas9-VP64 (Addgene, 180263) was obtained from Addgene. Custom GFP-tagged plasmids encoding three sgRNAs targeting the putative NR4A2 enhancer region cRE138020 (VectorBuilder, Vector ID: VB260221-1040tuh) or three non-targeting scramble gRNAs (VB260221-1038pmk) were obtained from VectorBuilder, where a full plasmid design is provided in **Figure S9I**. All plasmids were transformed in DH5α E. coli, followed by selection of individual colonies, expansion, and plasmid isolation as previously described. Lentivirus was generated in HEK293T in 10-cm dish as previously described with Lipofectamine 3000 and the same packaging plasmids. 24 hours after transfection, all media was removed and fresh complete RPMI with antibiotics and 1x ViralBoost Reagent (Alstem, VB100) was added. Lentiviral medium was collected 48 and 72 hours after transfection, and fresh complete RPMI was added after the first collection. The collected lentiviral medium was pooled and filtered through a sterile 0.45μm polyethersulfone (PES) filter. To concentrate the lentivirus, the filtered medium was added to the upper chamber of an Amicon Ultra Centrifugal filter, 10kDa molecular weight cut off, and spun at 3000 x g at 25 for 30 minutes. The lentivirus concentrate was resuspended to 1/10 the original volume with fresh RPMI. If the lentivirus was not immediately used it was flash frozen with liquid nitrogen prior to being stored at −80□. Jurkat cells were activated with Immunocult Human CD3/CD28 T Cell Activator (Stemcell, 10971) at a concentration of 25μL/mL. The next day, cells were diluted to a concentration of 2 million cells/mL and 2% volume per volume (v/v) concentrated mCherry-2A-dCas9-VP64 lentivirus was added with 5% v/v concentrated sgRNA-GFP lentivirus being added the following day. Three days after activation, fresh medium was added to bring cells to 7x105 cells/mL. Five days after activation, cells that were double positive for GFP and mCherry were sorted by a FACSymphony S6 Cell Sorter (BD Biosciences) to be plated as single cells in a 96-well tissue-culture plate. After 1 week of expansion, clones were run again to identify those with the highest GFP/mCherry expression and total RNA was isolated from these clones using the same process as described above. Total RNA was then reverse transcribed to cDNA using Lunascript RT or used to prepare sequencing libraries for bulk RNA-seq, as described below. QRT-PCR was run to quantify NR4A2 expression in the targeting and non-targeting conditions, where β-actin was used as a housekeeping gene for normalization.

### Bulk RNA Sequencing

RNA concentration and purity were evaluated using a Nanodrop 2000 (Thermo Fisher Scientific, ND2000). mRNA enrichment was accomplished with NEBNext Poly(A) mRNA Magnetic Isolation Module (NEB, E7490) immediately followed by library preparation using NEBNext Ultra II Directional RNA Library Prep Kit for Illumina (NEB, E7760) following the standard protocol according to the manufacturer’s instructions. Library concentration was measured with the Qubit dsDNA Quantification kit (Invitrogen, Q32851) using a SpectraMax M5 Multi-Mode Microplate Reader according to the manufacturer’s instructions. Fragment size and DNA quality were assessed using a 5300 Fragment Analyzer (Agilent) according to the manufacturer’s instruction after which all libraries were pooled in equimolar amounts for sequencing. Paired end sequencing was performed on an Illumina NovaSeq X Plus using a PE150-25B flow cell, 8bp dual indices, a 1% PhiX spike-in, and a 151 x 8 x 8 x 151 cycle configuration.

### Bulk RNA sequencing Data Analysis

Libraries were demultiplexed with bcl-convert and paired-end reads were aligned to hg38 with STAR 2.7.11b (GENCODE v46 index, sjdbOverhang 150), keeping unmapped reads. Gene-level counts were obtained with STAR --quantMode GeneCounts (reverse-stranded) and summed per gene symbol. Cell-line identity was confirmed by correlating expression-inferred copy-number profiles with CCLE references (37). Sample KO identity was confirmed by counting reads containing an exact match to each guide’s 20-nt protospacer followed by the first four scaffold bases (GTTT), in either orientation, normalized to reads per million unmapped reads. Genes passing edgeR filterByExpr in every cell line were retained, and expression was Trimmed Mean of M-values (TMM)-normalized log2 Counts Per Million (edgeR normalization), computed per cell line. To assess replicate agreement, gene expression (genes with mean CP10K ≥ 0.1 only) in KO samples per cell line was mean centered, and Spearman correlations between same KO replicate and different KO sample pairs were compared (one-sided Wilcoxon rank-sum test, FDR adjusted across cell lines). For proliferation analysis a proliferation score (using Seurat CellCycleScoring function) was computed per sample and scaled per cell line as (sample score − NTC mean)/SD. For each KO, the median across lines of the replicate means was compared with the NTC scores (one-sided Wilcoxon rank-sum test). To compare with the NCI-H2052 CROP-seq data, cell-cycle effects were regressed out of both datasets, KO profiles were then defined as the mean of all KOs minus each KO, and bulk profiles were averaged over lines. Spearman correlations were then computed across the 96 genes shared between the 500 most variable CROP-seq genes (Seurat vst) and the 500 most variable bulk genes. Same-KO pairs correlations were compared with different-KO pairs correlations by a one-sided Wilcoxon rank-sum test. Differential expression between cRE138020 enhancer CRISPRa and control samples was tested with DESeq2 (v1.46.0). The design was ∼ cell-cycle score + condition, with each sample’s cell-cycle score included as a covariate. For gene set enrichment analysis, all genes were ranked by the DESeq2 Wald statistic and analyzed by preranked GSEA with GSEApy (v1.3.1) against the MSigDB Hallmark 2020 and GO Biological Process 2023 gene set libraries, with FDR < 0.05 considered significant.

### Data availability

ScATAC-seq, scRNA-seq, CROP-seq, bulk RNA-seq, and 10x Visium data have been deposited to Gene Expression Omnibus (GEO) data repository under accession number GSE311510.

### Code availability

Custom scripts used for data preprocessing, analysis, and figure generation are publicly available at https://github.com/TsankovLab/PM_scATAC. Analyses were performed primarily using established single-cell analysis frameworks, including ArchR, Signac, and Seurat, together with additional R and Python scripts, as described in the **Methods** section.

## Acknowledgments

We thank all members of the Tsankov lab for their feedback as well as the ISMMS Genomics Core and Human Immune Monitoring Core for single cell profiling of PM tumor and CROP-seq samples. This work was supported by the American Association for Thoracic Surgery/Women in Thoracic Surgery Mid-career Investigator Grant to A.W. and Department of Army grant HT9425-23-1-0496 to A.M.T. This work was also supported in part through the computational and data resources and staff expertise provided by Scientific Computing and Data at the Icahn School of Medicine at Mount Sinai and supported by the Clinical and Translational Science Awards (CTSA) grant UL1TR004419 from the National Center for Advancing Translational Sciences. Research reported in this publication was also supported by the Office of Research Infrastructure of the National Institutes of Health under award number S10OD030463. The content is solely the responsibility of the authors and does not necessarily represent the official views of the National Institutes of Health.

## Author Contributions

B.G. and A.M.T. conceived and designed the study. B.G. led and performed the computational analysis with analytical assistance from W.L. and feedback from A.M.T. L.C. performed the CRISPR experiments with advice and resources from Y.S., S.S, B.H., and A.H., under the supervision of A.T., M.W, and A.M.T. K.D. and A.W. coordinated clinical tissue acquisition and processing. B.G. and A.M.T. wrote the manuscript with input and approval from all authors.

---

**Figure S1 supporting Figure 1.**
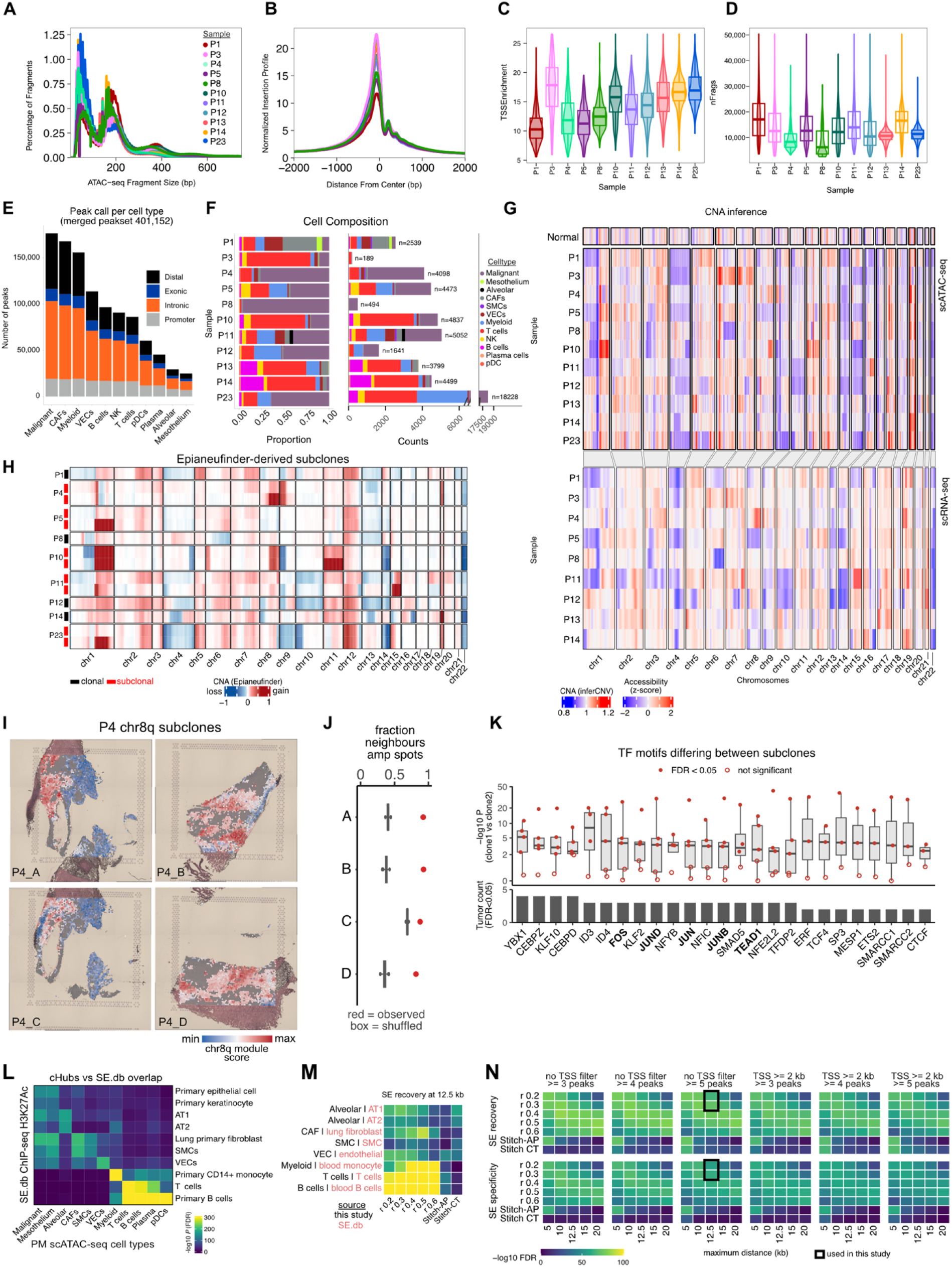
Single-cell atlas of chromatin accessibility in PM. **A,** Distribution of fragments length color-coded by sample. **B,** Distribution of fragments at transcript start site (TSS) loci color coded by sample. **C,** Distribution of TSS Enrichment scores split by sample. **D,** Distribution of total number of fragments split by sample. **E,** Total peak detected sliced by peak type and cell type. **F,** Cell composition of the 11 scATAC-seq samples as proportions (left) and cell counts (right). **G,** Heatmap of CNA inference from scATAC-seq on aggregated malignant cells for each tumor sample plus normal mesothelium after aggregating fragments in 10Mb genomic windows (top panel). Heatmap of CNA inference using inferCNV from paired scRNA-seq samples (bottom panel). **H**, Heatmap showing the CNA profiles of putative malignant cell subclones, where subclonal structure was inferred in 5 of 9 tumors analyzed. **I**, Spatial transcriptomics (10x Visium) of four PM tumor sections from sample P4 overlaid with normalized module scores of all chr8q genes expression. All four sections demonstrate a spatially correlated amplification of chr8q gene expression at malignant cell containing spots (**Methods**), where non-malignant spots shown in grey. **J**, Permutation analysis evaluating spatial co-localization of chr8q high spots, where red points show the fraction of malignant neighbors of chr8q-high spots that are also chr8q-high (adjacent spots on the Visium grid). One-sided permutation test in all four sections. **K**, For each of the five tumors with two inferred subclones (P4, P5, P10, P11, P23), chromVAR motif activity was compared between the cells assigned to the two putative clones (two-sided Wilcoxon rank-sum test, FDR adjusted). Top: Box plot of −log10 *P* values for the top 25 TFs, where points represent individual tumors (filled, FDR < 0.05; open, not significant). Bottom: Bar plot showing the number of tumors in which the motif activity differs significantly between inferred clones. **L,** Enrichment analysis of cHubs regions detected per cell type against a set of SE detected from ChIP-seq H3K27Ac data from the SE.db database (**Methods**). *P* values are reported as -log10 (hypergeometric test, FDR adjusted). **M,** Enrichment of SE sets (SE.db) in cell type-specific hubs for eight pairs of cHub cell type with matching SE set. CHubs were called by co-accessibility at correlation cut-offs r = 0.2–0.6, or by stitching peaks of the consensus (Stitch-AP) or cell-type-specific (Stitch-CT) peak sets with 12.5 kb maximum distance, ≥ 5 peaks per hub and no TSS exclusion. **N,** Top: SE recovery computed as mean of the matched pairs in panel **M.** Bottom: SE specificity computed as SE recovery minus enrichment of the same cell types for non-matching SE sets for each stitching method (rows) and maximum distance (columns), across all combinations of TSS exclusion and minimum peak number. The black framed panels show the setting used in this study. Values are capped at the 99th percentile. Box plot horizontal lines show the 25th, 50th (median) and 75th percentiles, with vertical whiskers extending to a maximum distance of 1.5× the interquartile range from the hinge. Data beyond the whisker ends are plotted individually. *P* values > 0.05 are not shown.

**Figure S2 supporting Figure 1.**
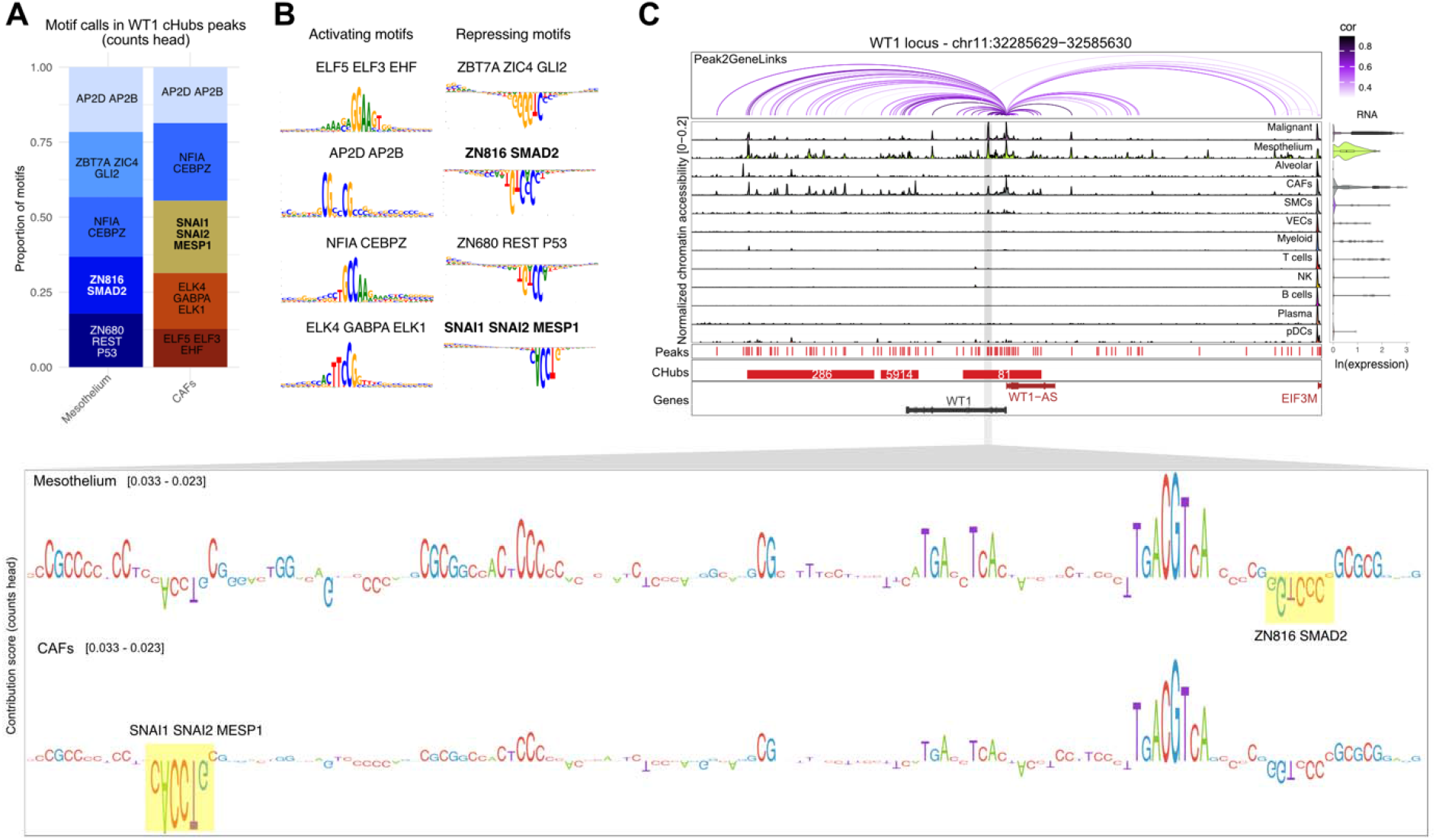
ChromBPNet deconvolves contribution of TF motifs at WT1 locus. **A,** Most frequent motifs calls detected from chromBPNet counts heads after training models on mesothelial and CAFs. **B,** *De-novo* motifs identified by chromBPNet matching top known motifs. Inverted motifs represent negative contributions to chromatin accessibility. C, Zoomed in peak region showing contribution scores (counts heads) from chromBPNet models trained on mesothelial cells and CAFs. Enriched motifs in either cell types are highlighted.

**Figure S3 supporting Figure 2.**
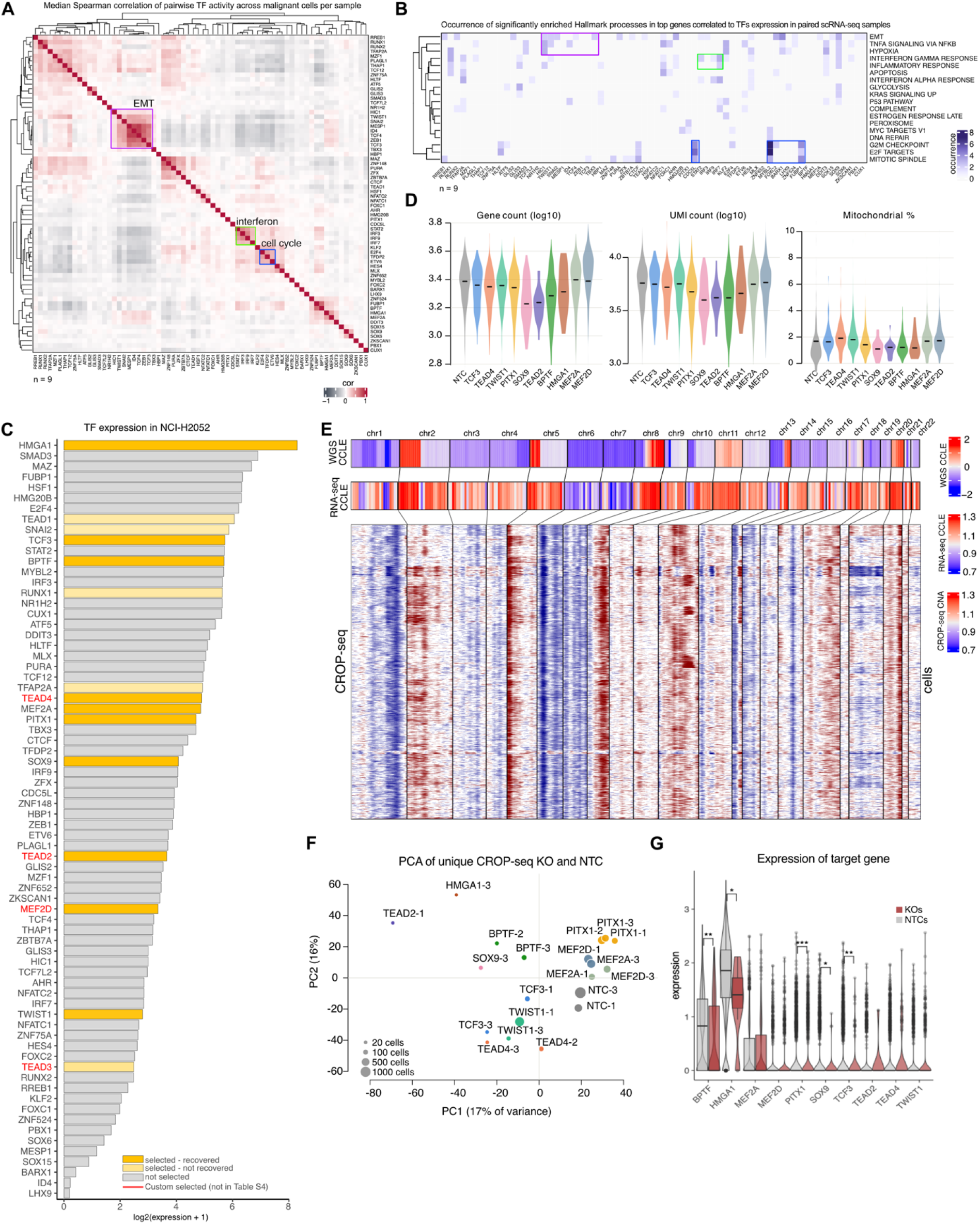
CRISPR droplet sequencing of computationally predicted pro-tumor TFs. **A,** Heatmap of TF co-activity using Spearman correlation showing the 68 TFs with higher activity and RNA expression in tumor cells as compared to normal mesothelium. Highlighted are clusters of TFs involved in the same biological pathway. **B,** Pathway enrichment analysis using HALLMARK database from MsigDB (accessed on 09/07/2025) of the top 100 correlated genes to TFs using scRNA-seq data from paired samples. Enrichments were carried out per samples and occurrence of significant pathway enrichment in each sample is shown. **C,** List of the 68 putative pro-tumorigenic TFs sorted by expression from bulk RNA-seq of NCI-H2052 mesothelioma cell line obtained from DepMap project data portal (https://depmap.org/portal) (86). **D,** QC plots of CROP-seq data showing gene counts, UMI counts and mitochondrial genes percentages. **E,** CNA inference on CROP-seq data using inferCNV (bottom heatmap) and comparison with CNA patterns derived from bulk RNA-seq (middle heatmap) and whole-exome sequencing data (top heatmap) of same cell line obtained from DepMap project data portal (https://depmap.org/portal) (86). **F,** PCA plot of mean transcriptomes averaged across cells (minimum 10 cells) uniquely mapping to guides shows clustering by target gene or NTCs. **G,** Distributions of target TF expression in cells with corresponding guides targeting that TF compared to NTC cells. Wilcoxon Rank Sum Test, FDR corrected. Box plot horizontal lines show the 25th, 50th (median) and 75th percentiles, with vertical whiskers extending to a maximum distance of 1.5× the interquartile range from the hinge. Data beyond the whisker ends are plotted individually. *\* P* value < 0.05, *\*\* P* value < 0.01, *\*\*\* P* value < 0.001, where *P* values > 0.05 are not shown.

**Figure S4 supporting Figure 2.**
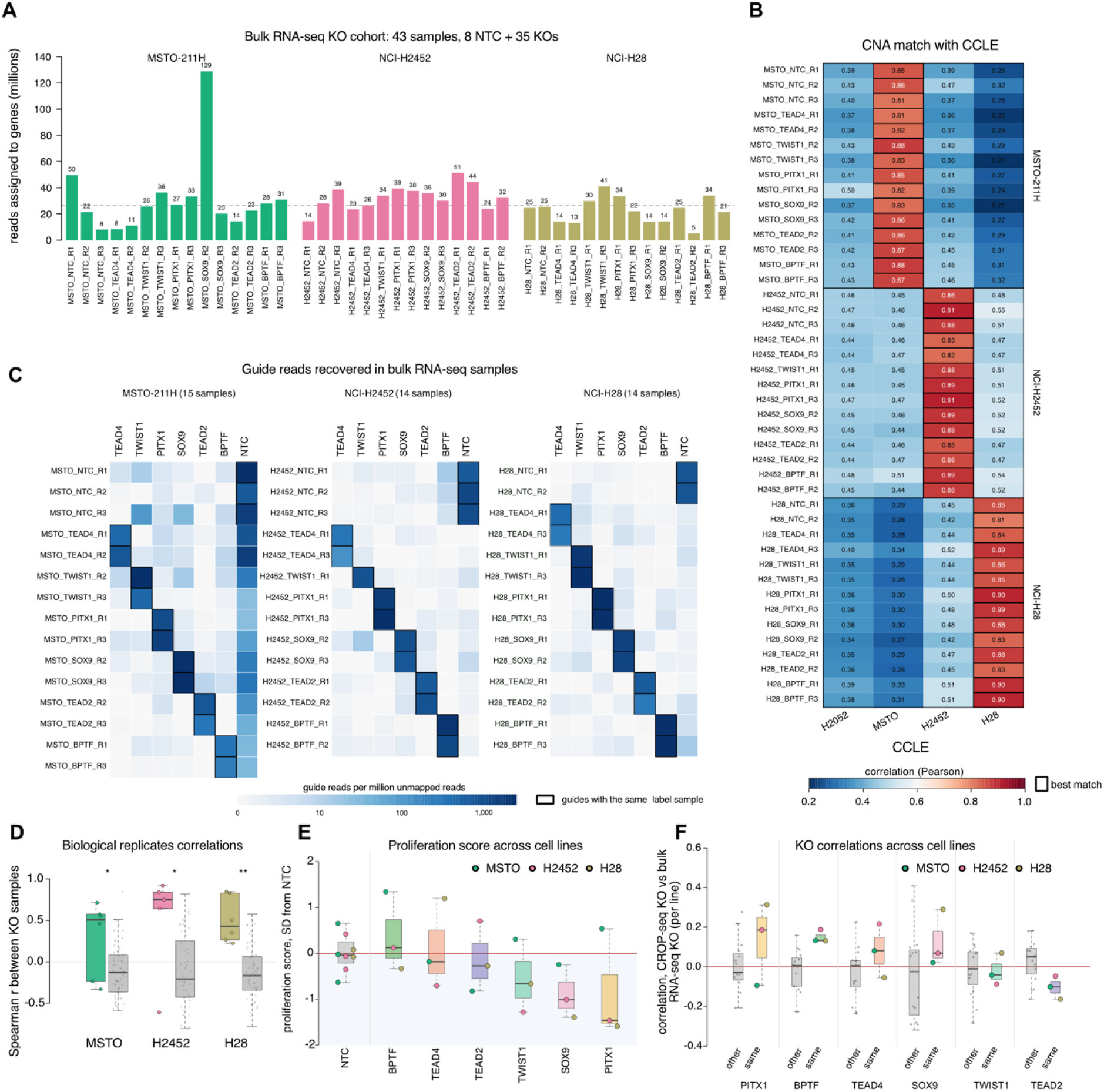
Bulk RNA-seq of TF KOs across mesothelioma cell lines. **A,** Breakdown of CRISPR KO and bulk RNA-seq experiment, which consisted of 35 KO and 8 NTC samples that passed QC. Bar plots show number of mapped reads. **B,** Heatmap showing the Pearson correlation between inferred CNA profile (**Methods**) for each bulk RNA-seq sample (rows) and the inferred CNA profiles for the four PM cell lines used in this study, using published bulk RNA-seq data from CCLE (37). All 43 samples showed highest correlation with the corresponding CCLE cell line data, confirming the expected cell line identity. **C,** Heatmap showing for each RNA-seq sample (rows) the recovered gRNA reads per target gene (columns) among the unmapped reads that contain a guide’s 20-nt protospacer followed by the scaffold sequence. Counts are normalized per million unmapped reads. **D,** Distribution of pairwise Spearman correlations between replicate samples targeting the same gene (colors) and between non-replicate samples (grey) across all expressed genes per cell line. FDR-corrected Wilcoxon Rank Sum Test. **E,** Proliferation score of KOs relative to NTCs normalized as the standard deviation (SD) of KO sample scores from the mean NTC proliferation score per cell line, where KO replicates were averaged (**Methods**). Circles, NTC and KO samples colored by cell line; red line, mean NTC level. **F**, Spearman correlation between CROP-seq KO pseudobulk expression profiles and bulk RNA-seq KO expression profiles comparing correlation between corresponding KO pairs (colored) versus unmatched KO pairs (grey), where KO replicates were averaged. Box plot horizontal lines show the 25th, 50th (median) and 75th percentiles, with vertical whiskers extending to a maximum distance of 1.5× the interquartile range from the hinge. Data beyond the whisker ends are plotted individually. *\*P* value < 0.05; *\*\*P* value < 0.01*; ***P* value < 0.001; *P* values > 0.05 are not shown.

**Figure S5 supporting Figure 3.**
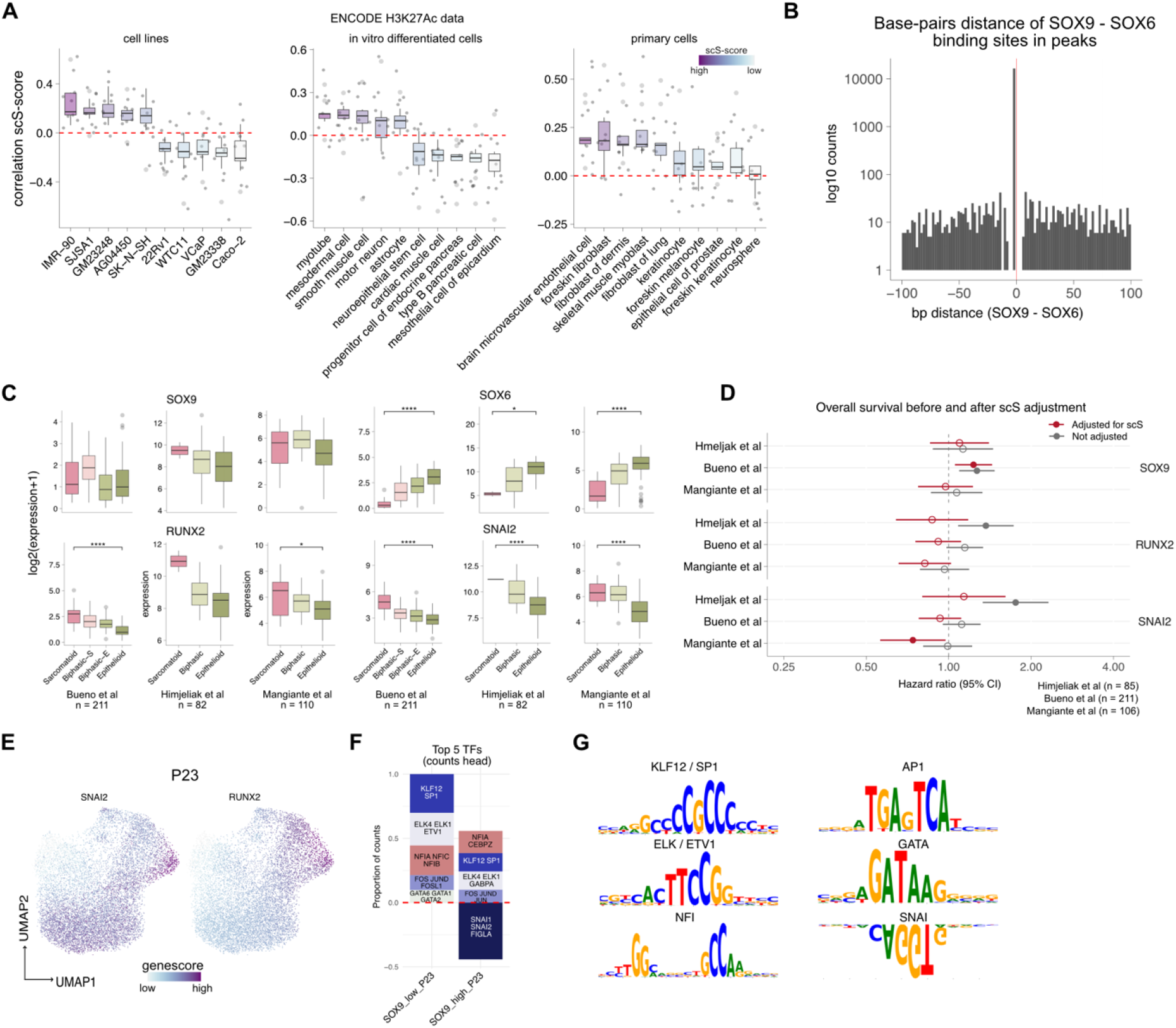
SOX9, RUNX2, and SNAI2 activity associates with sarcomatoid-like neoplastic state. **A,** Sample-level distributions of Spearman correlations between malignant cells’ scS-score and accessibility of open chromatin sets of peaks for a set of cell line, *in vitro* differentiated cells and primary cells (left, middle and right) profiled with ChIP-seq of H3K27Ac histone mark obtained from the ENCODE project (https://www.encodeproject.org/). **B,** Distribution of base-pair distances between the SOX9 and SOX6 binding sites (using the cisBP motif database (82)) within peaks in malignant cells. **C,** SOX9, RUNX2, SNAI2 and SOX6 log2 expression distributions across bulk RNA-seq samples from three large PM cohorts grouped by histological/molecular subtypes (16–18). Number of samples with subtype information is reported per cohort. **D,** Hazard ratios (HRs) and 95% confidence intervals from Cox proportional hazards models for overall survival, estimated per one-standard-deviation increase in gene expression in each of the three bulk RNA-seq PM cohorts (16–18). Number of samples with survival information is reported per cohort. Red indicates estimates adjusted for the mean scS score, while grey indicates unadjusted estimates. Gene expression and scS scores were standardized within each cohort (z-scores). Filled points indicate P < 0.05 (Wald test), dashed line marks HR = 1, and HR > 1 indicates an association between higher gene expression and worse overall survival. **E,** UMAPs of P23 cells with overlaid color-gradient representing genescore of SNAI2 and RUNX2. **F,** Most frequent motif calls detected from chromBPNet counts heads after training models on SOX9 high cells and SOX9 low cells. Motifs with negative proportions are associated with negative contributions to chromatin accessibility. **G,** *De-novo* motifs identified by chromBPNet matching top known motifs. Inverted motifs represent negative contributions to chromatin accessibility. Box plot horizontal lines show the 25th, 50th (median) and 75th percentiles, with vertical whiskers extending to a maximum distance of 1.5× the interquartile range from the hinge. Data beyond the whisker ends are plotted individually. *\* P* value < 0.05, *\*\* P* value < 0.01, *\*\*\* P* value < 0.001. where *P* values > 0.05 are not shown.

**Figure S6 supporting Figure 4.**
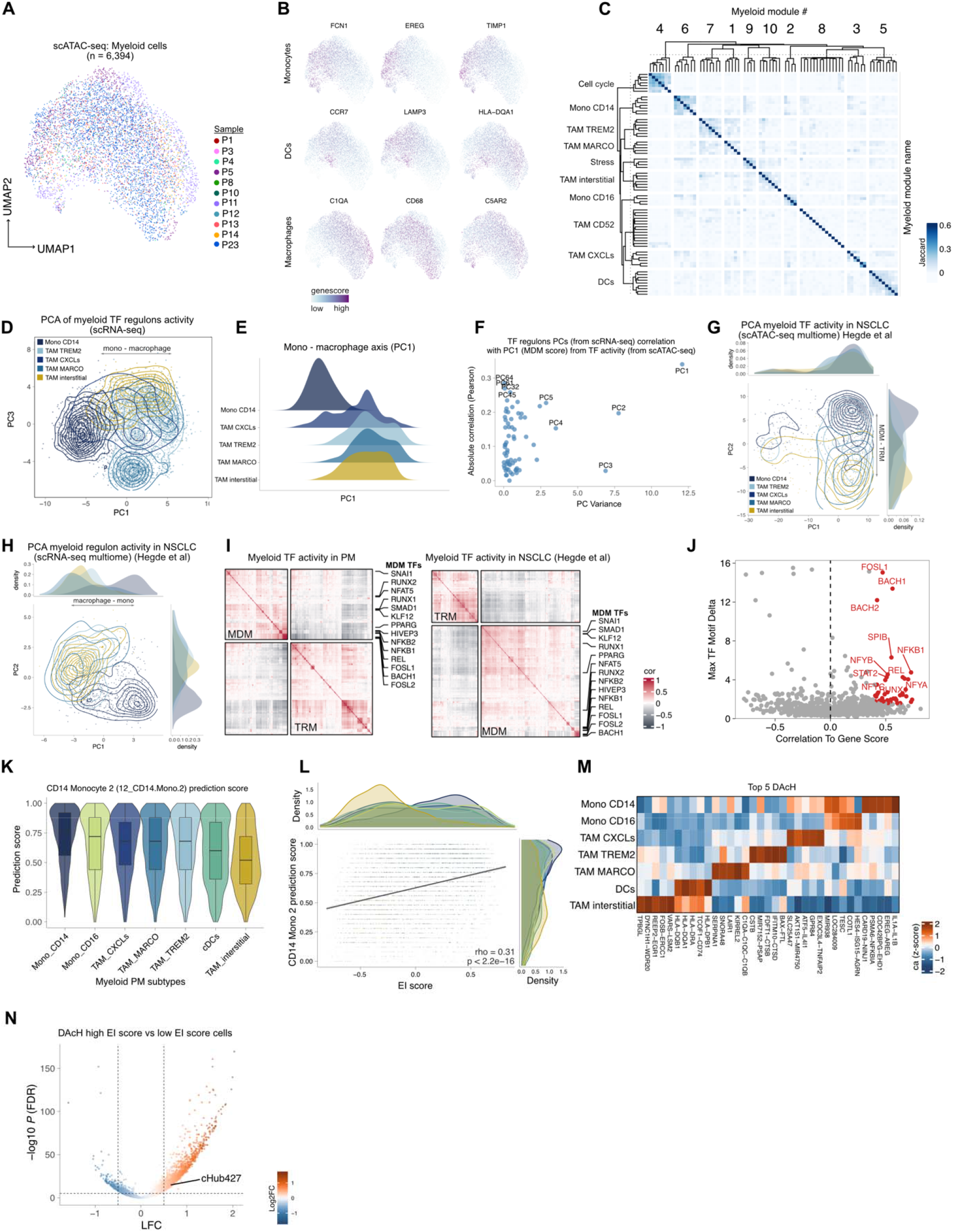
Epigenetic-based regulatory dichotomy captures putative macrophage ontogeny. **A**, UMAP of myeloid cell distribution color coded by sample. **B,** UMAPs of myeloid cell distributions with overlaid color-gradient representing genescore of *bona fide* marker genes of monocytes, DCs and macrophages. **C,** Heatmap of myeloid modules gene overlaps identified per sample from scRNA-seq paired data (**Methods**). **D,** PCA of Mono CD14 and TAM subsets on TF regulons inferred using SCENIC from paired scRNA-seq. Contours highlight areas of myeloid subsets. PC1 captured monocyte – macrophage regulatory axis. Since PC2 captured cell cycle variation, PC3 was shown instead. **E,** Cell density distributions on PC1 grouped by myeloid subset. **F,** Distribution of TF regulon PCs variance vs their correlation with PC1 from TF activity. **G,** PCAs of Mono CD14 and TAM subsets on TF activity. PC2 captured MDM – TRM regulatory axis. **H,** PCAs of Mono CD14 and TAM subsets on TF regulons inferred using SCENIC from paired scRNA-seq. PC1 captured macrophage – monocytes regulatory axis. Contours highlight areas of myeloid subsets. Derived from multiome data in NSCLC tumors (10). **I,** Heatmap of pairwise Spearman correlation between TF activities in PM (left) and NSCLC (right) myeloid. Heatmaps split into two anti-correlated TF regulatory modules. **J,** Distribution of maximum TF motif activity difference across myeloid clusters (Max TF Motif Delta, y-axis) versus relative genescore correlation (x-axis). TFs in red are considered as positive regulators of gene expression. **K,** Prediction scores derived from mapping CD14 Monocyte 2 scATAC-seq annotations from healthy individuals’ PBMCs (52) onto our PM-infiltrated myeloid cell subsets (columns, **Methods**). **L**, Scatterplot of CD14 Monocyte 2 prediction scores (y-axis) versus EI score (x-axis) across PM myeloid cells shows significant Spearman correlation. Density distributions for each myeloid subset are shown on top and right side. **M,** Heatmap of top DAcH across myeloid subsets. The number assigned to each cHub indicates its region size rank (rank=1 is largest), where cHub detection was recomputed within myeloid cells. **N,** Volcano plot DAcH between high and low EI score cells, highlighting differentially accessed cHub427 shown in Figure 4K. Box plot horizontal lines show the 25th, 50th (median) and 75th percentiles, with vertical whiskers extending to a maximum distance of 1.5× the interquartile range from the hinge. Data beyond the whisker ends are plotted individually. *P* values > 0.05 are not shown.

**Figure S7 supporting Figure 5.**
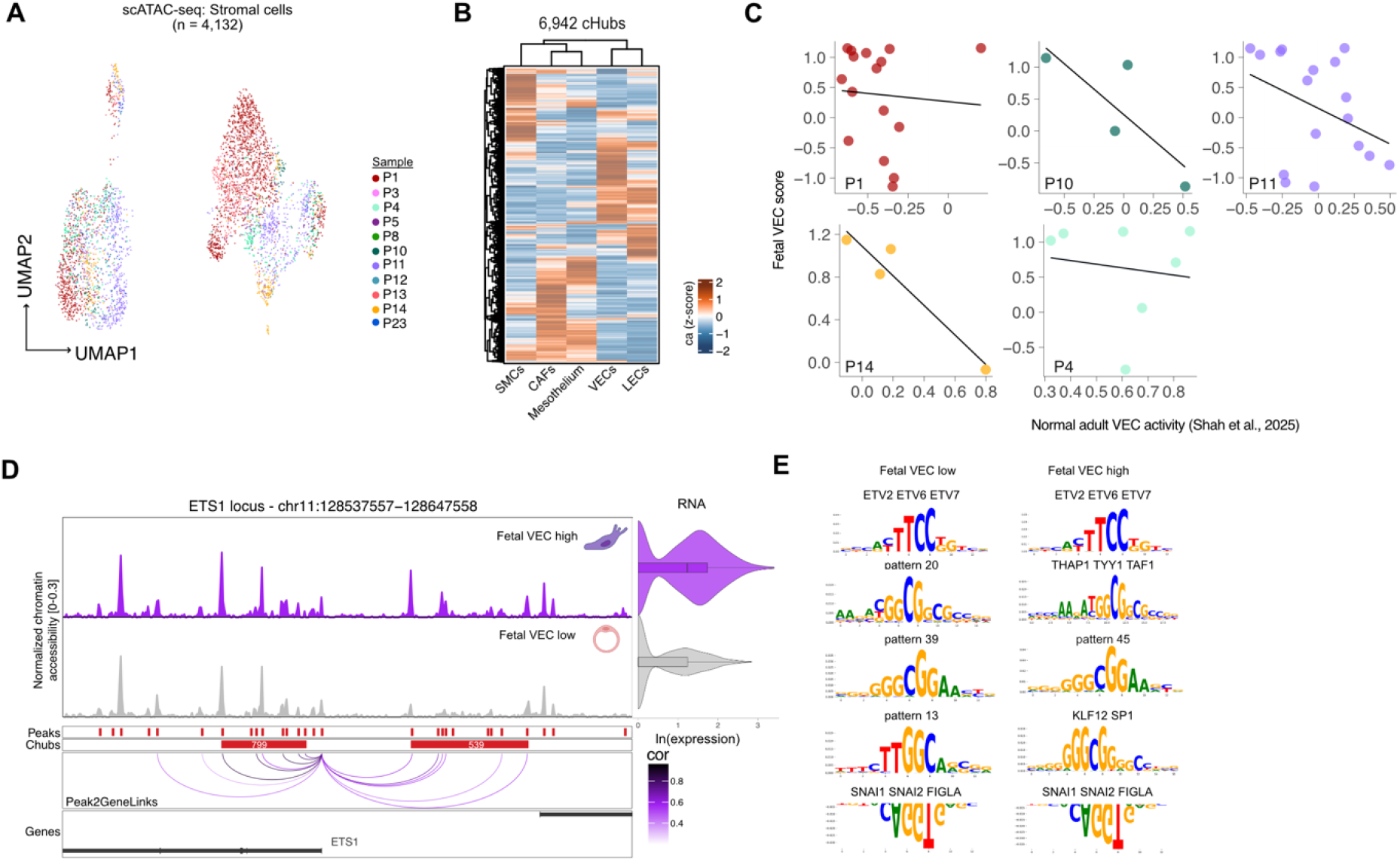
SNAI1 activity and cHub is associated with fetal-like endothelial cells in PM. **A,** UMAP of stromal cells color-coded by sample. **B,** Heatmap of cHubs detected across the stromal compartment. **C,** Per-sample distributions of metacells, with fetal VECs score on the y-axis and normal adult VECs activity on the x-axis. **D,** ETS1 locus with pseudobulk tracks of high and low fetal VECs score cells. Also displayed are ETS1’s RNA expression (violin plots, right), peak to gene correlation (purple arches, bottom), and cHubs detected (red bars). The number assigned to each cHub indicates its region size rank (rank=1 is largest), where cHub detection was recomputed within stromal cells. **E,** Top motifs identified by chromBPNet in VECs with high and low fetal VECs scores. Inverted motifs represent negative contributions to chromatin accessibility. Box plot horizontal lines show the 25th, 50th (median) and 75th percentiles, with vertical whiskers extending to a maximum distance of 1.5× the interquartile range from the hinge. Data beyond the whisker ends are plotted individually. *P* values > 0.05 are not shown.

**Figure S8 supporting Figure 6.**
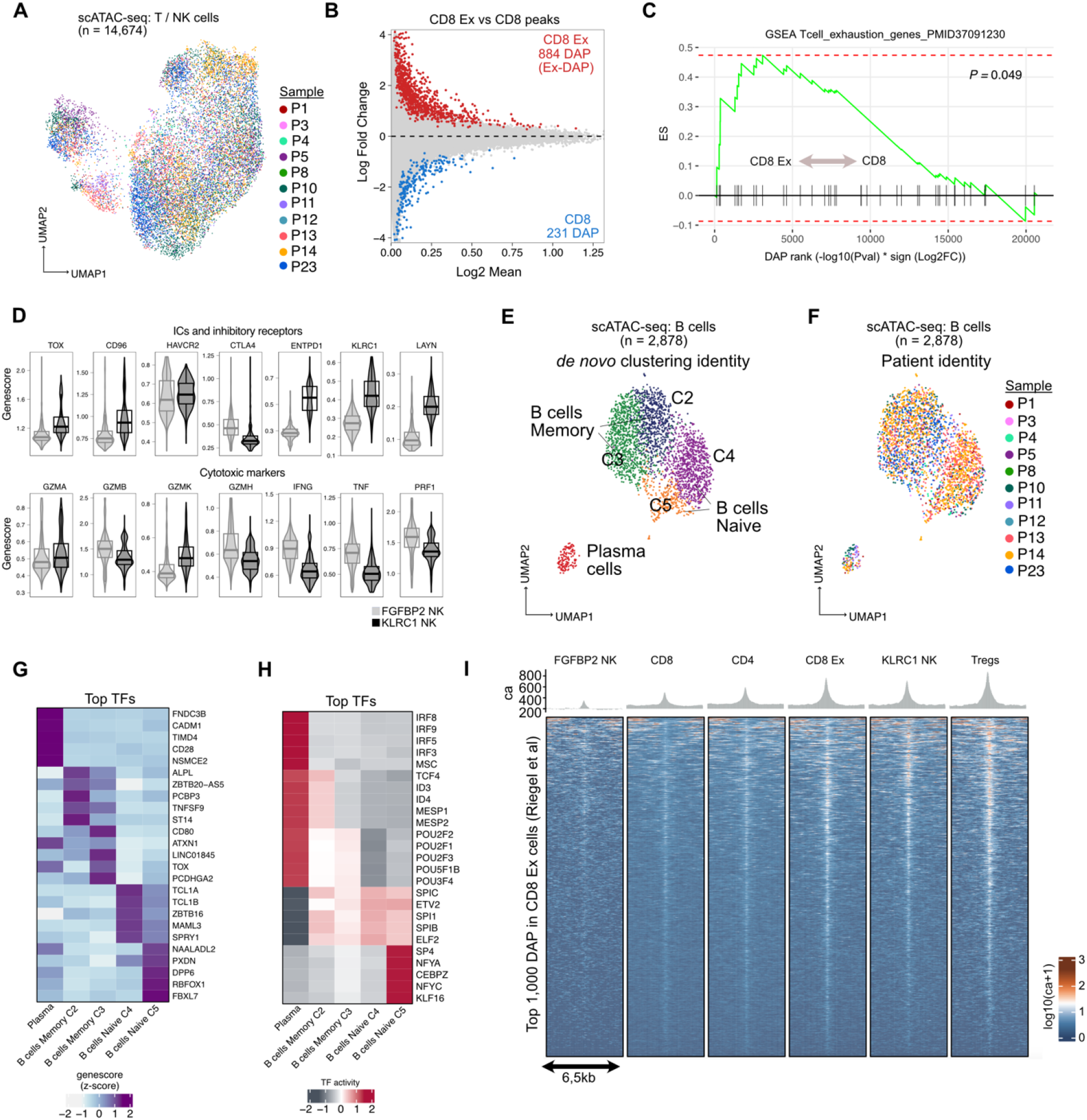
Exhausted CD8 T and KLRC1 NK cells share open chromatin regulatory circuits. **A,** UMAP of T-NK cells color-coded by sample. **B,** MA plot of DAP between CD8 Ex and CD8 cells. **C,** GSEA analysis on the ranked Ex-DAPs using a set of T-cell exhaustion genes (59). **D,** Genescore distributions of exhaustion markers, immune checkpoint molecules, and inhibitory receptors (top panel) as well as cytotoxic markers (bottom panel) split by NK cell subsets. **E,** UMAP of B and plasma cells color-coded by Louvain community detection clusters. **F,** UMAP of B and plasma cells color-coded by sample identity. **G,** Heatmap of top 5 genescores for each B and plasma cell cluster. **H,** Heatmap of top 5 TF activity for each B and plasma cell cluster. **I,** Chromatin accessibility of PM T-NK cell subsets at top 1,000 DAPs identified in CD8 Ex cells derived from four human cancer types (66). Box plot horizontal lines show the 25th, 50th (median) and 75th percentiles, with vertical whiskers extending to a maximum distance of 1.5× the interquartile range from the hinge. Data beyond the whisker ends are plotted individually. *P* values > 0.05 are not shown.

**Figure S9 supporting Figure 7.**
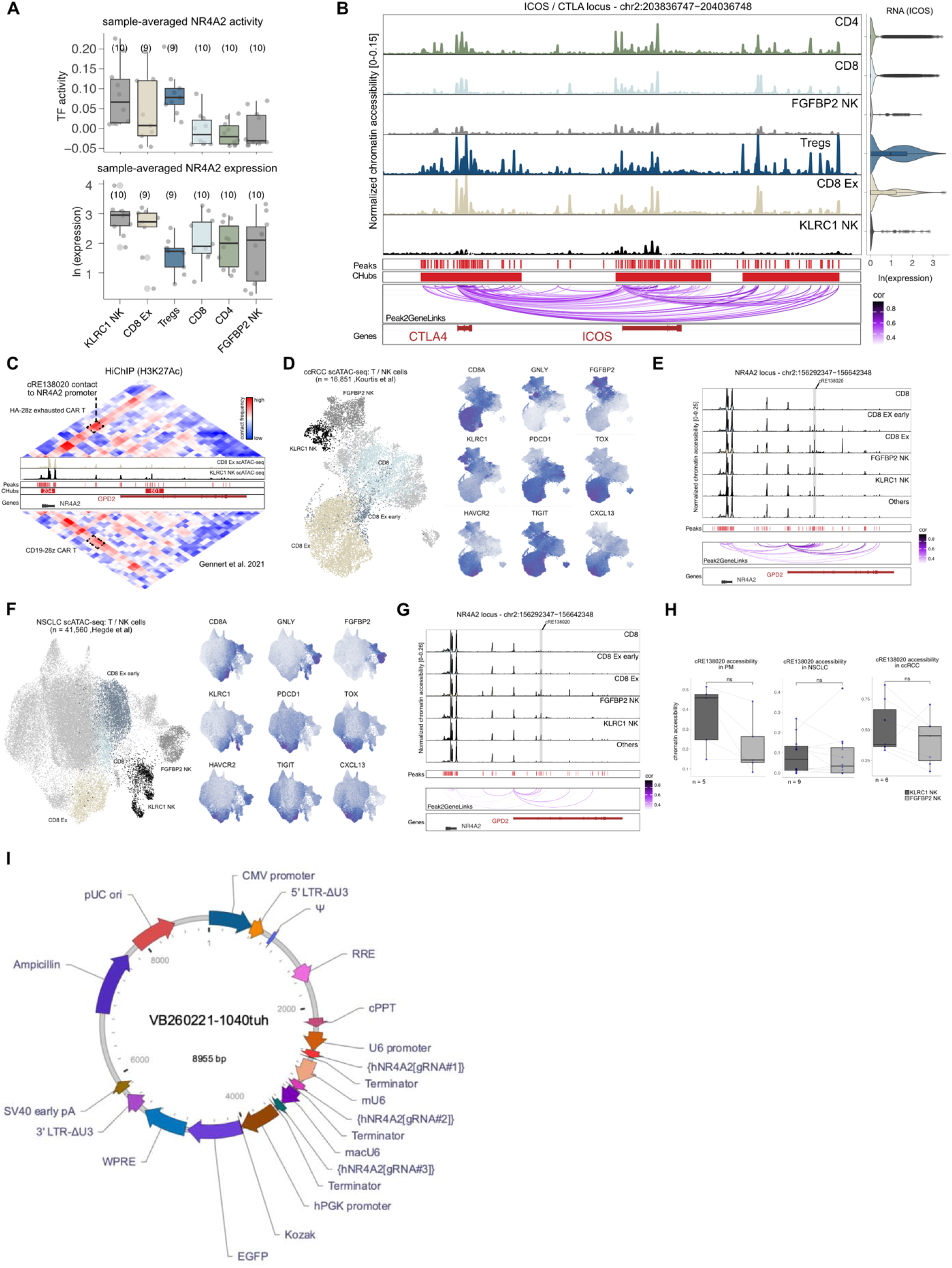
Exhaustion-specific enhancer cRE138020 likely regulates NR4A2. **A,** Sample distributions of NR4A2 activity and RNA expression in paired samples split by T/NK cell subsets. **B,** ICOS locus with pseudo-bulk tracks of T/NK cell subsets. Also displayed are ICOS’s RNA expression (violin plots, right), peak to gene correlation (purple arches, bottom), and cHubs detected (red bars). The number assigned to each cHub indicates its region size rank (rank=1 is largest), where cHub detection was recomputed within T/NK cells. **C,** Hi-ChIP data of HA-28z exhausted CAR T (top), as represented in Figure 7D, with comparative Hi-ChIP data of CD19-28z CAR T cells (bottom), which do not undergo exhaustion in *in vitro* culture (67). **D-E,** scATAC-seq data from ccRCC tumors (68) showing UMAP of T-NK cells color coded by cell subset (left), *bona fide* markers used to annotate cell subsets (right), and chromatin accessibility at the NR4A2 locus with the cRE138020 region highlighted and peak to gene correlation (purple arches, bottom) (**E)**. **F-G,** scATAC-seq data from NSCLC tumors (10) showing UMAP of T-NK cells color coded by cell subset (left), *bona fide* markers used to annotate cell subsets (right), and chromatin accessibility at the NR4A2 locus with the cRE138020 region highlighted and peak to gene correlation (purple arches, bottom) (**G)**. **H,** Sample distributions of chromatin accessibility at cRE138020 between KLRC1 NKs vs FGFBP2 NKs in our dataset (PM) in NSCLC (10) and in ccRCC (68). Student t-test. Ns = not significant. **I,** Vector map of the cRE138020-targeting plasmid (VB260221-1040tuh), showing annotated plasmid features and their arrangement within the 8,955-bp construct. Box plot horizontal lines show the 25th, 50th (median) and 75th percentiles, with vertical whiskers extending to a maximum distance of 1.5× the interquartile range from the hinge. Data beyond the whisker ends are plotted individually. *P* values > 0.05 are not shown.

## REFERENCES

1. Bxer E, Feigin N, Tschernichovsky R, Darnell NG, Greenwald AR, Hoefflin R, et al. Emerging clinical applications of single-cell RNA sequencing in oncology. Nat Rev Clin Oncol 2025;22(5):315–26 doi 10.1038/s41571-025-01003-3.

2. LaFave LM, Kartha VK, Ma S, Meli K, Del Priore I, Lareau C, et al. Epigenomic State Transitions Characterize Tumor Progression in Mouse Lung Adenocarcinoma. Cancer Cell 2020;38(2):212–28 e13 doi 10.1016/j.ccell.2020.06.006.

3. Becker WR, Nevins SA, Chen DC, Chiu R, Horning AM, Guha TK, et al. Single-cell analyses define a continuum of cell state and composition changes in the malignant transformation of polyps to colorectal cancer. Nat Genet 2022;54(7):985–95 doi 10.1038/s41588-022-01088-x.

4. Iglesia MD, Jayasinghe RG, Chen S, Terekhanova NV, Herndon JM, Storrs E, et al. Differential chromatin accessibility and transcriptional dynamics define breast cancer subtypes and their lineages. Nat Cancer 2024;5(11):1713–36 doi 10.1038/s43018-024-00773-6.

5. Poos AM, Prokoph N, Przybilla MJ, Mallm JP, Steiger S, Seufert I, et al. Resolving therapy resistance mechanisms in multiple myeloma by multiomics subclone analysis. Blood 2023;142(19):1633–46 doi 10.1182/blood.2023019758.

6. Regner MJ, Wisniewska K, Garcia-Recio S, Thennavan A, Mendez-Giraldez R, Malladi VS, et al. A multi-omic single-cell landscape of human gynecologic malignancies. Mol Cell 2021;81(23):4924–41 e10 doi 10.1016/j.molcel.2021.10.013.

7. Sundaram L, Kumar A, Zatzman M, Salcedo A, Ravindra N, Shams S, et al. Single-cell chromatin accessibility reveals malignant regulatory programs in primary human cancers. Science 2024;385(6713):eadk9217 doi 10.1126/science.adk9217.

8. Satpathy AT, Granja JM, Yost KE, Qi Y, Meschi F, McDermott GP, et al. Massively parallel single-cell chromatin landscapes of human immune cell development and intratumoral T cell exhaustion. Nat Biotechnol 2019;37(8):925–36 doi 10.1038/s41587-019-0206-z.

9. Gok Yavuz B, Khanmammadova N, Majeed Z, Ali MIH, Hasanov M, Bilen MA, et al. Mapping heterogeneity in the tumor microenvironment of renal cell carcinoma through single-cell omics. Trends Cancer 2025 doi 10.1016/j.trecan.2025.11.001.

10. Hegde S, Giotti B, Soong BY, Halasz L, Le Berichel J, Schaefer MM, et al. Myeloid progenitor dysregulation fuels immunosuppressive macrophages in tumours. Nature 2025;646(8087):1214–22 doi 10.1038/s41586-025-09493-y.

11. Abdel-Hakeem MS, Manne S, Beltra JC, Stelekati E, Chen Z, Nzingha K, et al. Epigenetic scarring of exhausted T cells hinders memory differentiation upon eliminating chronic antigenic stimulation. Nat Immunol 2021;22(8):1008–19 doi 10.1038/s41590-021-00975-5.

12. Weiss SA, Huang AY, Fung ME, Martinez D, Chen ACY, LaSalle TJ, et al. Epigenetic tuning of PD-1 expression improves exhausted T cell function and viral control. Nat Immunol 2024;25(10):1871–83 doi 10.1038/s41590-024-01961-3.

13. van Zandwijk N, Rasko JEJ, George AM, Frank AL, Reid G. The silent malignant mesothelioma epidemic: a call to action. Lancet Oncol 2022;23(10):1245–8 doi 10.1016/S1470-2045(22)00269-8.

14. Janes SM, Alrifai D, Fennell DA. Perspectives on the Treatment of Malignant Pleural Mesothelioma. N Engl J Med 2021;385(13):1207–18 doi 10.1056/NEJMra1912719.

15. Baas P, Scherpereel A, Nowak AK, Fujimoto N, Peters S, Tsao AS, et al. First-line nivolumab plus ipilimumab in unresectable malignant pleural mesothelioma (CheckMate 743): a multicentre, randomised, open-label, phase 3 trial. Lancet 2021;397(10272):375–86 doi 10.1016/S0140-6736(20)32714-8.

16. Bueno R, Stawiski EW, Goldstein LD, Durinck S, De Rienzo A, Modrusan Z, et al. Comprehensive genomic analysis of malignant pleural mesothelioma identifies recurrent mutations, gene fusions and splicing alterations. Nat Genet 2016;48(4):407–16 doi 10.1038/ng.3520.

17. Hmeljak J, Sanchez-Vega F, Hoadley KA, Shih J, Stewart C, Heiman D, et al. Integrative Molecular Characterization of Malignant Pleural Mesothelioma. Cancer Discov 2018;8(12):1548–65 doi 10.1158/2159-8290.CD-18-0804.

18. Mangiante L, Alcala N, Sexton-Oates A, Di Genova A, Gonzalez-Perez A, Khandekar A, et al. Multiomic analysis of malignant pleural mesothelioma identifies molecular axes and specialized tumor profiles driving intertumor heterogeneity. Nat Genet 2023;55(4):607–18 doi 10.1038/s41588-023-01321-1.

19. Blum Y, Meiller C, Quetel L, Elarouci N, Ayadi M, Tashtanbaeva D, et al. Dissecting heterogeneity in malignant pleural mesothelioma through histo-molecular gradients for clinical applications. Nature Communications 2019;10 doi ARTN 1333 10.1038/s41467-019-09307-6.

20. Severson DT, Freyaldenhoven S, Wadowski B, Hung YP, Hughes T, Yeap BY, et al. Multiomic, Histologic, and scRNA-seq Profiling of Pleural Mesothelioma Reveals Negative Prognosis Associated With a Novel Uncommitted Molecular Phenotype. J Thorac Oncol 2025 doi 10.1016/j.jtho.2025.11.019.

21. Giotti B, Dolasia K, Zhao W, Cai P, Sweeney R, Merritt E, et al. Single-Cell View of Tumor Microenvironment Gradients in Pleural Mesothelioma. Cancer Discov 2024;14(11):2262–78 doi 10.1158/2159-8290.CD-23-0017.

22. Schep AN, Wu B, Buenrostro JD, Greenleaf WJ. chromVAR: inferring transcription-factor-associated accessibility from single-cell epigenomic data. Nat Methods 2017;14(10):975–8 doi 10.1038/nmeth.4401.

23. Demos C, Johnson J, Andueza A, Park C, Kim Y, Villa-Roel N, et al. Sox13 is a novel flow-sensitive transcription factor that prevents inflammation by repressing chemokine expression in endothelial cells. Front Cardiovasc Med 2022;9 doi ARTN 979745 10.3389/fcvm.2022.979745.

24. Zakrzewska A, Cui C, Stockhammer OW, Benard EL, Spaink HP, Meijer AH. Macrophage-specific gene functions in Spi1-directed innate immunity. Blood 2010;116(3):E1–E11 doi DOI 10.1182/blood-2010-01-262873.

25. Hirai H, Yang B, Garcia-Barrio MT, Rom O, Ma PX, Zhang J, Chen YE. Direct Reprogramming of Fibroblasts Into Smooth Muscle-Like Cells With Defined Transcription Factors-Brief Report. Arterioscler Thromb Vasc Biol 2018;38(9):2191–7 doi 10.1161/ATVBAHA.118.310870.

26. Akao K, Sato T, Mishiro-Sato E, Mukai S, Ghani FI, Kondo-Ida L, et al. TEAD-independent cell growth of Hippo-inactive mesothelioma cells: Unveiling resistance to TEAD inhibitor K-975 through MYC signaling activation. Mol Cancer Ther 2024 doi 10.1158/1535-7163.MCT-24-0308.

27. Ramakrishnan A, Symeonidi A, Hanel P, Schmid KT, Richter ML, Schubert M, Colome-Tatche M. epiAneufinder identifies copy number alterations from single-cell ATAC-seq data. Nat Commun 2023;14(1):5846 doi 10.1038/s41467-023-41076-1.

28. Hnisz D, Abraham BJ, Lee TI, Lau A, Saint-Andre V, Sigova AA, et al. Super-enhancers in the control of cell identity and disease. Cell 2013;155(4):934–47 doi 10.1016/j.cell.2013.09.053.

29. Whyte WA, Orlando DA, Hnisz D, Abraham BJ, Lin CY, Kagey MH, et al. Master transcription factors and mediator establish super-enhancers at key cell identity genes. Cell 2013;153(2):307–19 doi 10.1016/j.cell.2013.03.035.

30. Tsankov AM, Gu H, Akopian V, Ziller MJ, Donaghey J, Amit I, et al. Transcription factor binding dynamics during human ES cell differentiation. Nature 2015;518(7539):344–9 doi 10.1038/nature14233.

31. Pliner HA, Packer JS, McFaline-Figueroa JL, Cusanovich DA, Daza RM, Aghamirzaie D, et al. Cicero Predicts cis-Regulatory DNA Interactions from Single-Cell Chromatin Accessibility Data. Mol Cell 2018;71(5):858–71 e8 doi 10.1016/j.molcel.2018.06.044.

32. Wang Y, Song C, Zhao J, Zhang Y, Zhao X, Feng C, et al. SEdb 2.0: a comprehensive super-enhancer database of human and mouse. Nucleic Acids Res 2023;51(D1):D280–D90 doi 10.1093/nar/gkac968.

33. Park S, Schalling M, Bernard A, Maheswaran S, Shipley GC, Roberts D, et al. The Wilms tumour gene WT1 is expressed in murine mesoderm-derived tissues and mutated in a human mesothelioma. Nat Genet 1993;4(4):415–20 doi 10.1038/ng0893-415.

34. Pampari A, Shcherbina A, Kvon EZ, Kosicki M, Nair S, Kundu S, et al. ChromBPNet: bias factorized, base-resolution deep learning models of chromatin accessibility reveal cis-regulatory sequence syntax, transcription factor footprints and regulatory variants. bioRxiv 2025 doi 10.1101/2024.12.25.630221.

35. Shah VS, Waghray A, Lin B, Bhagwat A, Monga I, Slyper M, et al. Single cell profiling of human airway identifies tuft-ionocyte progenitor cells displaying cytokine-dependent differentiation bias in vitro. Nat Commun 2025;16(1):5180 doi 10.1038/s41467-025-60441-w.

36. Datlinger P, Rendeiro AF, Schmidl C, Krausgruber T, Traxler P, Klughammer J, et al. Pooled CRISPR screening with single-cell transcriptome readout. Nat Methods 2017;14(3):297–301 doi 10.1038/nmeth.4177.

37. Barretina J, Caponigro G, Stransky N, Venkatesan K, Margolin AA, Kim S, et al. The Cancer Cell Line Encyclopedia enables predictive modelling of anticancer drug sensitivity. Nature 2012;483(7391):603–7 doi 10.1038/nature11003.

38. Yang MH, Wu KJ. TWIST activation by hypoxia inducible factor-1 (HIF-1): implications in metastasis and development. Cell Cycle 2008;7(14):2090–6 doi 10.4161/cc.7.14.6324.

39. Cao X, Pfaff SL, Gage FH. YAP regulates neural progenitor cell number via the TEA domain transcription factor. Genes Dev 2008;22(23):3320–34 doi 10.1101/gad.1726608.

40. Liu XL, Zuo R, Ou WB. The hippo pathway provides novel insights into lung cancer and mesothelioma treatment. J Cancer Res Clin Oncol 2018;144(11):2097–106 doi 10.1007/s00432-018-2727-0.

41. Zeng ZX, Gu SS, Ouardaoui N, Tymm C, Yang L, Wong CJ, et al. Hippo Signaling Pathway Regulates Cancer Cell-Intrinsic MHC-II Expression. Cancer Immunol Res 2022;10(12):1559–69 doi 10.1158/2326-6066.Cir-22-0227.

42. Mansfield AS, Symanowski JT, Peikert T. Systematic review of response rates of sarcomatoid malignant pleural mesotheliomas in clinical trials. Lung Cancer 2014;86(2):133–6 doi 10.1016/j.lungcan.2014.08.017.

43. Yang YH, Gomez N, Infarinato N, Adam RC, Sribour M, Baek I, et al. The pioneer factor SOX9 competes for epigenetic factors to switch stem cell fates. Nature Cell Biology 2023;25(8):1185-+ doi 10.1038/s41556-023-01184-y.

44. Sakai D, Suzuki T, Osumi N, Wakamatsu Y. Cooperative action of Sox9, Snail2 and PKA signaling in early neural crest development. Development 2006;133(7):1323–33 doi 10.1242/dev.02297.

45. Fukui T, Sumitomo R, Menju T, Kobayashi M, Sakai H, Date H. M2-like tumor-associated macrophages may promote tumor progression in malignant pleural mesothelioma. Transl Oncol 2025;54:102324 doi 10.1016/j.tranon.2025.102324.

46. Lievense LA, Cornelissen R, Bezemer K, Kaijen-Lambers ME, Hegmans JP, Aerts JG. Pleural Effusion of Patients with Malignant Mesothelioma Induces Macrophage-Mediated T Cell Suppression. J Thorac Oncol 2016;11(10):1755–64 doi 10.1016/j.jtho.2016.06.021.

47. Leader AM, Grout JA, Maier BB, Nabet BY, Park MD, Tabachnikova A, et al. Single-cell analysis of human non-small cell lung cancer lesions refines tumor classification and patient stratification. Cancer Cell 2021;39(12):1594–609 e12 doi 10.1016/j.ccell.2021.10.009.

48. Tamura A, Hirai H, Yokota A, Kamio N, Sato A, Shoji T, et al. C/EBPβ is required for survival of Ly6C monocytes. Blood 2017;130(16):1809–18 doi 10.1182/blood-2017-03-772962.

49. Gallant S, Gilkeson G. ETS transcription factors and regulation of immunity. Arch Immunol Ther Exp (Warsz) 2006;54(3):149–63 doi 10.1007/s00005-006-0017-z.

50. DeKoter RP, Singh H. Regulation of B lymphocyte and macrophage development by graded expression of PU.1. Science 2000;288(5470):1439–41 doi 10.1126/science.288.5470.1439.

51. Ashburner M, Ball CA, Blake JA, Botstein D, Butler H, Cherry JM, et al. Gene Ontology: tool for the unification of biology. Nature Genetics 2000;25(1):25–9 doi Doi 10.1038/75556.

52. Granja JM, Klemm S, McGinnis LM, Kathiria AS, Mezger A, Corces MR, et al. Single-cell multiomic analysis identifies regulatory programs in mixed-phenotype acute leukemia. Nat Biotechnol 2019;37(12):1458–65 doi 10.1038/s41587-019-0332-7.

53. Xu MS, Zhang T, Xia RL, Wei YQ, Wei XW. Targeting the tumor stroma for cancer therapy. Mol Cancer 2022;21(1) doi ARTN 208 10.1186/s12943-022-01670-1.

54. Rojas A, De Val S, Heidt AB, Xu S-M, Bristow J, Black BL. Gata4 expression in lateral mesoderm is downstream of BMP4 and is activated directly by Forkhead and GATA transcription factors through a distal enhancer element. 2005.

55. Dubey S, Dubey PK, Umeshappa CS, Ghebre YT, Krishnamurthy P. Inhibition of RUNX1 blocks the differentiation of lung fibroblasts to myofibroblasts. J Cell Physiol 2022;237(4):2169–82 doi 10.1002/jcp.30684.

56. Zolak JS, Jagirdar R, Surolia R, Karki S, Oliva O, Hock T, et al. Pleural mesothelial cell differentiation and invasion in fibrogenic lung injury. Am J Pathol 2013;182(4):1239–47 doi 10.1016/j.ajpath.2012.12.030.

57. Lomeli H, Starling C, Gridley T. Epiblast-specific Snai1 deletion results in embryonic lethality due to multiple vascular defects. BMC Res Notes 2009;2:22 doi 10.1186/1756-0500-2-22.

58. Oh SY, Kim JY, Park C. The ETS Factor, ETV2: a Master Regulator for Vascular Endothelial Cell Development. Mol Cells 2015;38(12):1029–36 doi 10.14348/molcells.2015.0331.

59. Zhang C, Sheng Q, Zhang X, Xu K, Jin X, Zhou W, et al. Prioritizing exhausted T cell marker genes highlights immune subtypes in pan-cancer. iScience 2023;26(4):106484 doi 10.1016/j.isci.2023.106484.

60. Netskar H, Pfefferle A, Goodridge JP, Sohlberg E, Dufva O, Teichmann SA, et al. Pan-cancer profiling of tumor-infiltrating natural killer cells through transcriptional reference mapping. Nature Immunology 2024;25(8) doi 10.1038/s41590-024-01884-z.

61. Serger C, Rebuffet L, Sandholzer MT, Rackwitz W, Fusi I, Herzig P, et al. Cytotoxic CD39 tumor-associated NK cells respond to NKG2A blockade in lung cancer. Sci Immunol 2026;11(120) doi ARTN eaeb6645 10.1126/sciimmunol.aeb6645.

62. Moroney JB, Vasudev A, Pertsemlidis A, Zan H, Casali P. Integrative transcriptome and chromatin landscape analysis reveals distinct epigenetic regulations in human memory B cells. Nat Commun 2020;11(1):5435 doi 10.1038/s41467-020-19242-6.

63. King HW, Wells KL, Shipony Z, Kathiria AS, Wagar LE, Lareau C, et al. Integrated single-cell transcriptomics and epigenomics reveals strong germinal center-associated etiology of autoimmune risk loci. Sci Immunol 2021;6(64) doi ARTN eabh3768 10.1126/sciimmunol.abh3768.

64. Itahashi K, Irie T, Yuda J, Kumagai S, Tanegashima T, Lin YT, et al. BATF epigenetically and transcriptionally controls the activation program of regulatory T cells in human tumors. Sci Immunol 2022;7(76) doi ARTN eabk0957 10.1126/sciimmunol.abk0957.

65. Gordon SM, Chaix J, Rupp LJ, Wu JM, Madera S, Sun JC, et al. The Transcription Factors T-bet and Eomes Control Key Checkpoints of Natural Killer Cell Maturation. Immunity 2012;36(1):55–67 doi 10.1016/j.immuni.2011.11.016.

66. Riegel D, Romero-Fernandez E, Simon M, Adenugba AR, Singer K, Mayr R, et al. Integrated single-cell profiling dissects cell-state-specific enhancer landscapes of human tumor-infiltrating CD8(+) T cells. Mol Cell 2023;83(4):622–36 e10 doi 10.1016/j.molcel.2022.12.029.

67. Gennert DG, Lynn RC, Granja JM, Weber EW, Mumbach MR, Zhao Y, et al. Dynamic chromatin regulatory landscape of human CAR T cell exhaustion. Proc Natl Acad Sci U S A 2021;118(30) doi 10.1073/pnas.2104758118.

68. Kourtis N, Wang Q, Wang B, Oswald E, Adler C, Cherravuru S, et al. A single-cell map of dynamic chromatin landscapes of immune cells in renal cell carcinoma. Nat Cancer 2022;3(7):885–98 doi 10.1038/s43018-022-00391-0.

69. Ma F, Ye H, He HH, Gerrin SJ, Chen S, Tanenbaum BA, et al. SOX9 drives WNT pathway activation in prostate cancer. J Clin Invest 2016;126(5):1745–58 doi 10.1172/JCI78815.

70. Ma Y, Shepherd J, Zhao D, Bollu LR, Tahaney WM, Hill J, et al. SOX9 Is Essential for Triple-Negative Breast Cancer Cell Survival and Metastasis. Mol Cancer Res 2020;18(12):1825–38 doi 10.1158/1541-7786.MCR-19-0311.

71. Santos JC, Carrasco-Garcia E, Garcia-Puga M, Aldaz P, Montes M, Fernandez-Reyes M, et al. SOX9 Elevation Acts with Canonical WNT Signaling to Drive Gastric Cancer Progression. Cancer Res 2016;76(22):6735–46 doi 10.1158/0008-5472.CAN-16-1120.

72. Lau CM, Adams NM, Geary CD, Weizman OE, Rapp M, Pritykin Y, et al. Epigenetic control of innate and adaptive immune memory. Nature Immunology 2018;19(9):963-+ doi 10.1038/s41590-018-0176-1.

73. Viel S, Vivier E, Walzer T, Marçais A. Targeting metabolic dysfunction of CD8 T cells and natural killer cells in cancer. Nat Rev Drug Discov 2025;24(3):190–208 doi 10.1038/s41573-024-01098-w.

74. Sabag B, Puthenveetil A, Levy M, Joseph N, Doniger T, Yaron O, et al. Dysfunctional natural killer cells can be reprogrammed to regain anti-tumor activity. Embo J 2024;43(13):2552–81 doi 10.1038/s44318-024-00094-5.

75. Chen J, Lopez-Moyado IF, Seo H, Lio CJ, Hempleman LJ, Sekiya T, et al. NR4A transcription factors limit CAR T cell function in solid tumours. Nature 2019;567(7749):530–4 doi 10.1038/s41586-019-0985-x.

76. Nyberg WA, Bernard PL, Ngo W, Wang CH, Ark J, Rothrock A, et al. In vivo site-specific engineering to reprogram T cells. Nature 2026 doi 10.1038/s41586-026-10235-x.

77. Stuart T, Srivastava A, Madad S, Lareau CA, Satija R. Single-cell chromatin state analysis with Signac. Nat Methods 2021;18(11):1333–41 doi 10.1038/s41592-021-01282-5.

78. Levine JH, Simonds EF, Bendall SC, Davis KL, Amir el AD, Tadmor MD, et al. Data-Driven Phenotypic Dissection of AML Reveals Progenitor-like Cells that Correlate with Prognosis. Cell 2015;162(1):184–97 doi 10.1016/j.cell.2015.05.047.

79. Stuart T, Butler A, Hoffman P, Hafemeister C, Papalexi E, Mauck WM, 3rd, et al. Comprehensive Integration of Single-Cell Data. Cell 2019;177(7):1888–902 e21 doi 10.1016/j.cell.2019.05.031.

80. Granja JM, Corces MR, Pierce SE, Bagdatli ST, Choudhry H, Chang HY, Greenleaf WJ. ArchR is a scalable software package for integrative single-cell chromatin accessibility analysis. Nat Genet 2021;53(3):403–11 doi 10.1038/s41588-021-00790-6.

81. Korsunsky I, Millard N, Fan J, Slowikowski K, Zhang F, Wei K, et al. Fast, sensitive and accurate integration of single-cell data with Harmony. Nat Methods 2019;16(12):1289–96 doi 10.1038/s41592-019-0619-0.

82. Weirauch MT, Yang A, Albu M, Cote AG, Montenegro-Montero A, Drewe P, et al. Determination and inference of eukaryotic transcription factor sequence specificity. Cell 2014;158(6):1431–43 doi 10.1016/j.cell.2014.08.009.

83. Morabito S, Reese F, Rahimzadeh N, Miyoshi E, Swarup V. hdWGCNA identifies co-expression networks in high-dimensional transcriptomics data. Cell Rep Methods 2023;3(6):100498 doi 10.1016/j.crmeth.2023.100498.

84. Aibar S, Gonzalez-Blas CB, Moerman T, Huynh-Thu VA, Imrichova H, Hulselmans G, et al. SCENIC: single-cell regulatory network inference and clustering. Nat Methods 2017;14(11):1083–6 doi 10.1038/nmeth.4463.

85. Patel AP, Tirosh I, Trombetta JJ, Shalek AK, Gillespie SM, Wakimoto H, et al. Single-cell RNA-seq highlights intratumoral heterogeneity in primary glioblastoma. Science 2014;344(6190):1396–401 doi 10.1126/science.1254257.

86. Arafeh R, Shibue T, Dempster JM, Hahn WC, Vazquez F. The present and future of the Cancer Dependency Map. Nat Rev Cancer 2025;25(1):59–73 doi 10.1038/s41568-024-00763-x.

